# The Calmodulin-like proteins, CML13 and CML14 Function as Myosin Light Chains for the Class XI Myosins in *Arabidopsis*

**DOI:** 10.1101/2024.07.11.603113

**Authors:** Kyle Symonds, Liam Duff, Vikas Dwivedi, Eduard Belausov, Lalita Pal, Motoki Tominaga, Takeshi Haraguchi, Einat Sadot, Kohji Ito, Wayne A Snedden

## Abstract

Myosins are a crucial motor protein associated with the actin cytoskeleton in eukaryotic cells. Structurally, myosins form heteromeric complexes, with smaller light chains such as calmodulin (CaM) bound to isoleucine–glutamine (IQ) domains in the neck region. These interactions facilitate mechano-enzymatic activity. Recently, we identified Arabidopsis CaM-like (CML) proteins CML13 and CML14 as interactors with proteins containing multiple IQ domains, that function as the myosin VIII light chains. This study demonstrates that CaM, CML13, and CML14 specifically bind to the neck region of all 13 Arabidopsis myosin XI isoforms, with some preference among the CaM/CML-IQ domains. Additionally, we observed distinct residue preferences within the IQ domains for CML13, CML14, and CaM. *In vitro* experiments revealed that recombinant CaM, CML13, and CML14 exhibit calcium-independent binding to the IQ domains of myosin XIs. Furthermore, when co-expressed with MAP65-1–myosin fusion proteins containing the IQ domains of myosin XIs, CaM, CML13, and CML14 co-localize to microtubules. *In vitro* actin motility assays demonstrated that recombinant CML13, CML14, and CaM function as myosin XI light chains. A *cml13* T-DNA mutant exhibited a shortened primary root phenotype that was complemented by the wild-type CML13 and was similar to that observed in a triple myosin XI mutant (*xi3KO*). Overall, our data indicate that Arabidopsis CML13 and CML14 are novel myosin XI light chains that likely participate in a breadth of myosin XI functions.

**Highlight:** Myosin XI proteins play a crucial role in the plant cytoskeleton, but their associated light chains have remained unidentified. Here, we show that calmodulin-like proteins, CML13 and CML14, serve as light chains for myosin XI, similar to their role for myosin VIII proteins

## Introduction

The cytoskeleton is a dynamic and intricate system that plays a crucial role in organizing the intracellular environment. Within the cytoskeleton, the acto-myosin network stands out as a ubiquitous system, consisting of myosin motor proteins bound to actin polymers. The myosin superfamily can be broadly categorized into two groups: conventional and unconventional.

Conventional myosins form filaments and primarily function in animal muscle contraction and cell motility (Altman, 2013). In contrast, unconventional myosins do not form filaments and participate in diverse processes, including organelle trafficking, cell and organism growth, and nuclear rearrangements (Hartman *et al*., 2011; Altman, 2013; Nebenführ and Dixit, 2018). These unconventional myosins consist of distinct domains: the motor, neck, and tail domains, and they may also include coiled-coil domains.

Among the 79 phylogenetic classes of the myosin superfamily (Kollmar and Mühlhausen, 2017) only two, VIII and XI, are present in plants (Nebenführ and Dixit, 2018). The Arabidopsis genome encodes four class VIII and 13 class XI myosins (Avisar *et al*., 2009; Peremyslov *et al*., 2010; Haraguchi *et al*., 2018). The roles of myosin VIIIs are unclear, and a report using a quadruple knockout line for all myosin VIIIs did not reveal strong phenotypes (Talts *et al*., 2016). Class VIII myosins have been implicated in processes such as plasmodesmatal transport, endocytosis, and hypocotyl elongation (Baluska *et al*., 2001; Golomb *et al*., 2008; Sattarzadeh *et al*., 2008; Symonds *et al*., 2024b). Their enzymatic profile suggests tension sensor and/or tension generator activity, rather than fast movement (Haraguchi *et al*., 2014; Henn and Sadot, 2014; Rula *et al*., 2018; Olatunji *et al*., 2023). Recent reports also highlight the involvement of myosin VIIIs and XIs in the movement of viral particles from tobacco mosaic, rice stripe, and rice grassy stunt viruses, as well as the VirE2 protein from *Agrobacterium tumefaciens* (Pitzalis and Heinlein, 2017; Liu *et al*., 2024). Myosin XI functions are well-studied and include various intracellular processes. Myosin XI-1, -2, and -K have emerged as the main players in the active movement of vesicles, organelles, the bulk flow of the cytoplasm (Peremyslov *et al*., 2010; Cai *et al*.. 2014). Indeed, the engineering of Arabidopsis myosin XI-2 with the faster *Chara braunii* myosin XI motor domain resulted in increased Arabidopsis and *Camelina sativa* growth owing to faster cytoplasmic transport and larger cell sizes (Tominaga *et al*., 2013; Duan *et al*., 2020). Further, myosin XIK and XI-1 where shown to play a role during cell division (Abu-Abied *et al*., 2018; Huang *et al*., 2024). Some of the other Arabidopsis myosin XI family members have specialized functions. For example, myosin XI-F primarily acts in the elongation and gravitropic response of the inflorescence stem (Okamoto *et al*., 2015). Myosin XI-B, -C, and -E are involved in the movement and delivery of vesicles in the apical growth of pollen tubes (Madison *et al*., 2015; Wang *et al*., 2020; Tian *et al*., 2021), myosin XI-A and -D mutants had reduced pollen tube growth and seed set rate but presented no apparent effect on vesicle movement (Madison *et al*., 2015), and myosin XI-I is solely involved in maintaining nuclear envelope shape and positioning (Tamura *et al*., 2013; Zhou *et al*., 2015). Interestingly, the broadly expressed myosin XI-H and the pollen-specific XI-J have currently unknown functions (Haraguchi *et al*., 2018). Unlike all other myosins studied to date, XI-G is reported to function in the active movement of F-actin filaments during sperm nuclear migration, rather than the canonical myosin role of cargo transporters (Ali *et al*., 2020). In barely (*Hordeum vulgare)*, the ortholog of myosin XI-A was identified as the causative mutation in *Ror1* (Required for *mlo* powdery mildew resistance) and was speculated to be involved in delivering antifungal and/or cell wall cargoes to the sites of pathogen recognition (Acevedo-Garcia *et al*., 2022). Ror1 could also be forming a signalosome or nanodomain complex with immunity signalling components, similar to Arabidopsis myosin XI-K forming a BIK1 (Botrytis-induced kinase 1) nanodomain to facilitate innate immunity (Wang *et al*., 2024). The ortholog of myosin XI-I in *Zea mays,* Opaque1, was necessary for proper asymmetric cell divisions during stomatal development (Nan *et al*., 2023). Myosin XIs have broad cellular and physiological functions, yet the biochemical regulation of their activity *in vitro* and *in vivo* is predominantly unstudied.

In a generalized model of myosin protein architecture, the catalytic motor domain of myosins interacts with actin filaments and is responsible for ATP hydrolysis. The neck domain binds one or more myosin light chains (MLCs) via IQ (isoleucine–glutamine) motifs. When present, the coiled-coil domain facilitates homodimerization, while the globular tail domain binds to various cargoes (Peremyslov *et al*., 2013; Kurth *et al*., 2017; Duan and Tominaga, 2018). Although our understanding of plant myosin neck domains remains limited, they serve as the lever arm during mobility, making them essential for proper myosin function. These necks contain one or more IQ domains, characterized by the consensus motif IQXXXRGXXXR, which are arranged in proximity within the primary sequence. These IQ domains act as the binding sites for MLCs, providing the rigidity necessary for myosin movement and the calcium regulation of the holoenzyme (Heissler and Sellers, 2014). Plant myosin VIIIs are calcium-independent motors with three or four IQ domains that can be occupied by calmodulin (CaM) and/or CaM-like (CML) proteins CML13 and CML14 as their MLCs, exhibiting some IQ specificity (Haraguchi *et al*., 2014; Symonds *et al*., 2024b). Current models suggest that CaM typically acts as the MLC on the IQ1 domain, closest to the motor domain, and undergoes conformational changes while remaining bound in the presence and absence of calcium (Symonds *et al*., 2024b). This conformational change likely contributes to the motor’s calcium inhibition during elevated calcium concentrations *in vivo* (Duan and Tominaga, 2018). CML13 and CML14 specifically interact with IQ2 and are required for maximal myosin motility *in vitro (Symonds et al., 2024b)*. However, their role in myosin VIII regulation remains unknown. IQ3 and, if present, IQ4 are non-specific IQs, serving primarily as structural support rather than functional regulation, and can be occupied by CaM, CML13, or CML14 both *in planta* and *in vitro (Symonds et al., 2024b)*. In contrast, myosin XIs typically have six IQ domains, with CaM currently being the only known MLC (Nebenführ and Dixit, 2018). Myosin XIs also likely have alternative MLCs to CaM as a previous report on the motility of myosin XI motor-neck domains found that CaM alone was insufficient for most myosin motilities *in vitro (Haraguchi et al., 2018)*. Given the crucial functions that myosin XIs perform in plant growth and development, understanding their MLCs would advance our knowledge of intracellular actin-based transport. Our present study found that CML13 and CML14, along with CaM, interacted with the neck domains of all myosin XIs and facilitated the motility of myosin XI-H, I, and K *in vitro*. Notably, CML13 and CML14 exhibited IQ specificity for IQ2 and IQ4 within the myosin neck domains, however, the basis for this specificity remains unknown. Finally, the partial knockout of *CML13*, *cml13-1*, phenocopied the reduced root length phenotype of the myosin XI-1, -2, and -K triple knockout, *xi3KO*. This phenotype was also rescuable by expressing CML13-GFP in the *cml13-1* background, *cml13R 1* and *2*. Taken together, our data indicate that CML13 and CML14 are novel MLCs important for myosin XIs activity and function.

## Results

### Sequence comparisons of Arabidopsis myosin XI neck domains

Our previous study on Arabidopsis myosin VIIIs found that CaM, CML13, and CML14 act as the MLCs for these myosins by binding to their neck domains. In comparison, the neck regions of the class XI myosins are unique from those of the myosin VIIIs, as they possess six tandem IQ motifs in contrast to three or four found in myosin VIIIs (Fig. 1A). In the myosin XI alignment, the main IQ consensus residues (IQXXXRGXXXR) are underlined and show that each myosin XI is predicted to contain six IQ motifs. The neck domain of myosin XI-G is four residues shorter than the other family members and likely has had two deletion events removing two residues each immediately following IQ3 and IQ4. The IQ motif sequences are rather variable, with only IQ2 possessing the five core IQ motif residues as the consensus. Myosin XI IQ1 has a conserved Ser/Thr (marked by asterisks) in place of the consensus Gly. Interestingly, a previous report suggested that the Thr in this position in myosin XI-I is a putative phosphosite (Mergner *et al*., 2020). However, whether other myosin XIs are also phosphorylated at this site and the physiological or biochemical consequences of phosphorylation are currently unknown. IQ5 was the least conserved of the group, containing only the first three of the five core residues with considerable variability within the rest of the motif. A phylogenetic tree (Fig. 1B) shows the subgrouping of the 13 myosin XI isoforms encoded in the Arabidopsis genome. IQ6 is the only IQ motif that does not possess the conserved Iso/Leu/Val at position one, instead, the consensus is a Thr in this position except for myosin XI-I and XI-J which conform to the canonical motif. Most myosins possess a relatively close paralog except for myosin XI-F, -I, and -J which are phylogenetically more distant from the other myosins.

**Figure 1.**
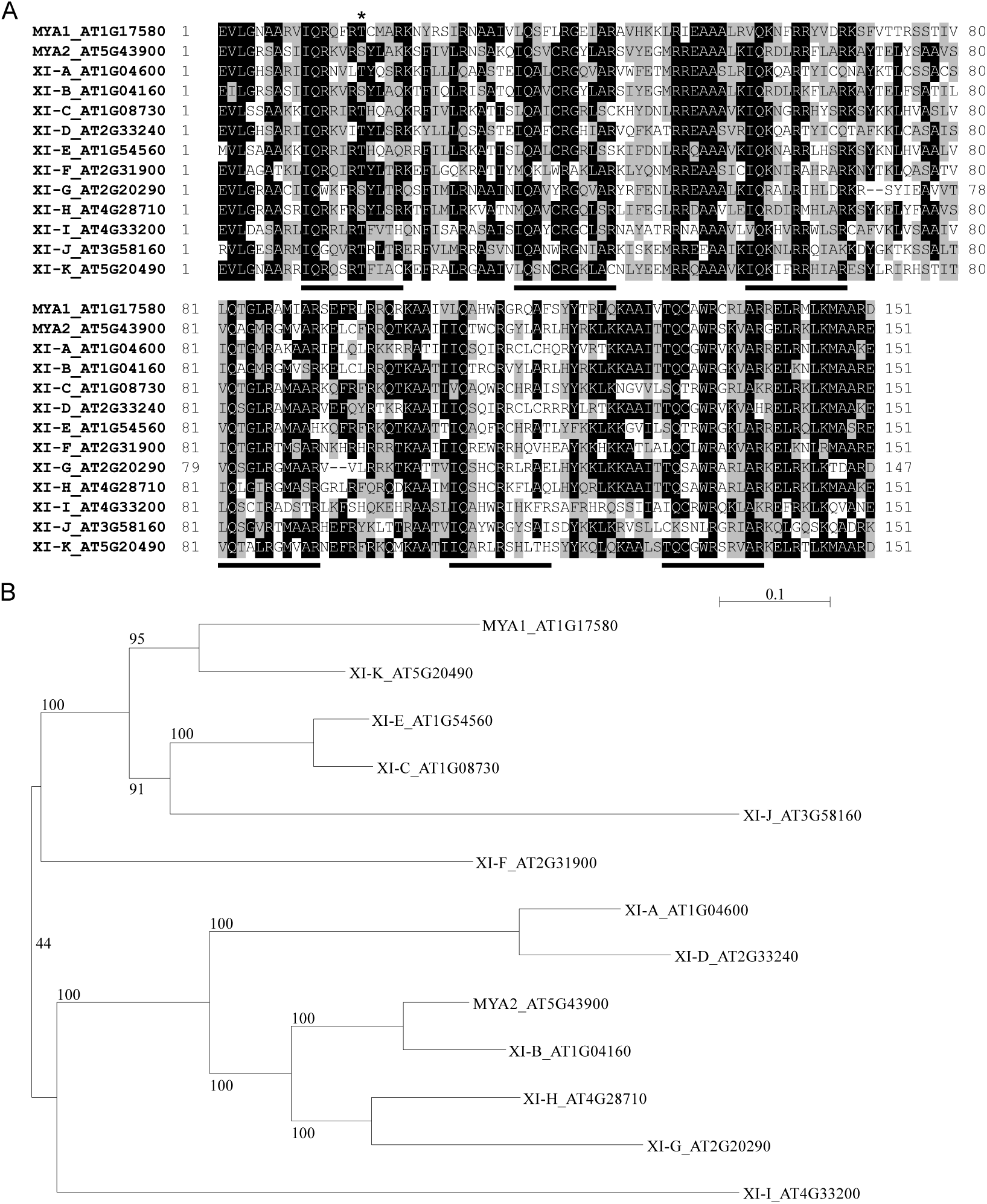
Protein sequence alignment of myosin XI neck domains (residues ∼750-900) and phylogenetic tree of full-length class XI myosins. A) amino acid residues were shaded black if they were greater than 50% homologous and progressively lighter shades of grey for lower identities. The consensus sequence of the predicted IQ motifs within the neck domains were underlined and the site of the putative myosin XI-I IQ1 phosphosite was denoted by an asterisk on top of the alignment. Clustal omega (Sievers and Higgins, 2014) was used to construct the alignment and BioEdit (Hall, 1999) was used to generate the image. B) A phylogenetic tree of class XI myosin full-length proteins was constructed using SeaView V5.0 and the PhyML algorithm with the Blosum62 scoring matrix and 1000 bootstrap replicates shown on the tree’s branches.

### CML13, CML14, and CaM Interact with Myosin XIs in planta

To assess the interaction of the myosin XI neck domains to the myosin VIII MLCs (CML13, CML14, and CaM) we utilized the *in planta* split luciferase (SL) protein-protein interaction assay. A schematic of the neck domain containing the six IQ motifs is shown in Fig. 2A. The neck domains of all 13 myosin XIs were cloned into the N-terminal Luciferase vector and tested for interaction with CML13, CML14, and CaM, with CML42 as the negative control. Fig 2B shows a heat map of the average log_2_ fold change of RLU produced in the SL normalized to CML42 with the same myosin XI neck domain. We observed that CML13, CML14, and CaM interacted with each of the 13 myosin XI neck domains. Most of the neck domains tested produced higher RLUs with CML13 and CML14 compared to CaM except for myosin XI-1, XI-2, and XI-G. Myosins XI-A and XI-I had the highest signal with CML13 and CML14 potentially implying tight binding to these neck domains.

**Figure 2.**
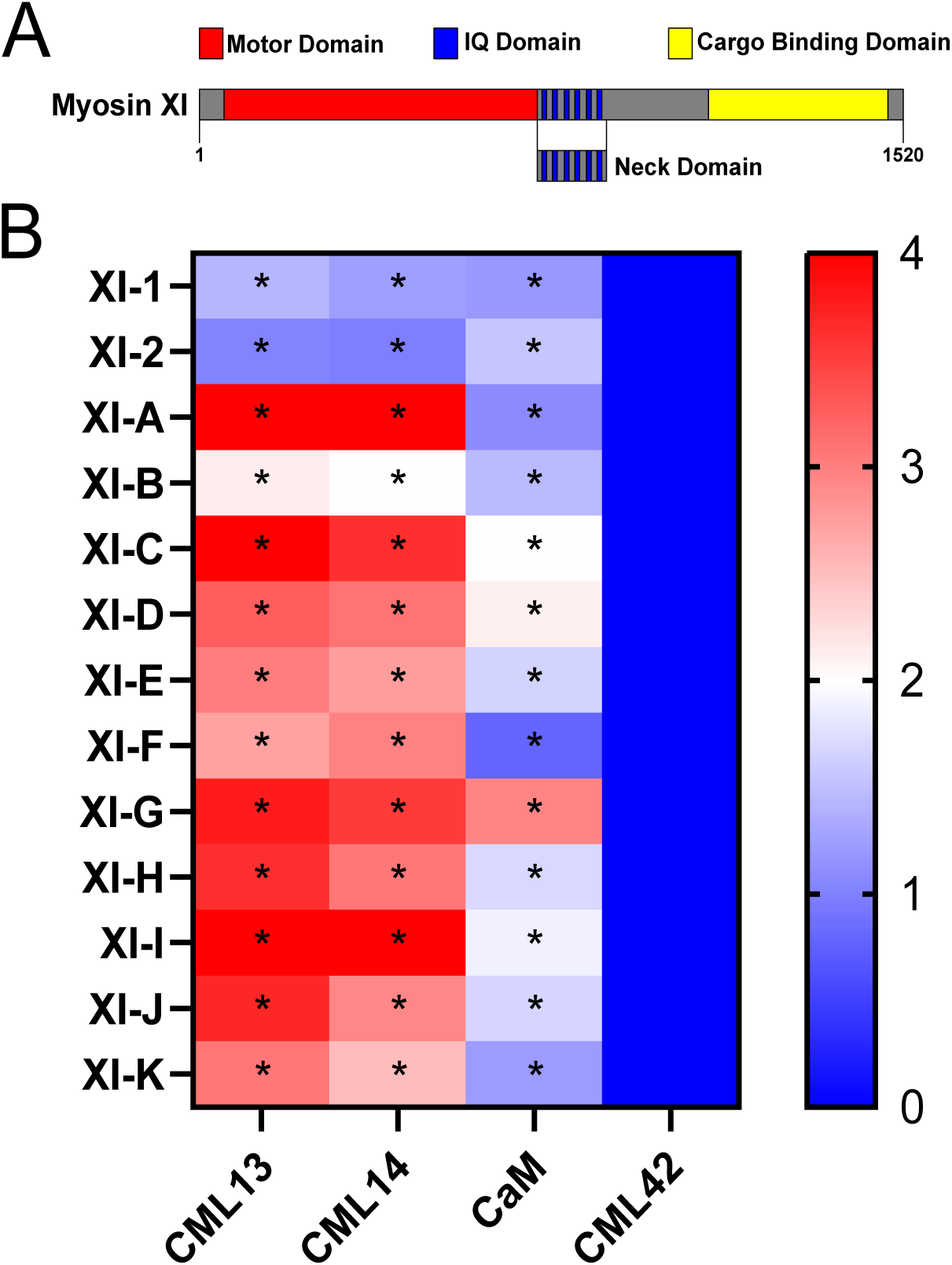
Split-luciferase protein interaction of Arabidopsis CMLs with the neck region of class XI myosins *in planta*. *N. benthamiana* leaves were co-infiltrated with *Agrobacterium* harboring an NLuc-myosin and CLuc-CML plasmid and were allowed to transiently express these fusion proteins for four days. A) Schematic of a typical myosin XI protein domain structure highlighting the neck domain to be tested in the SL assay. B) A heat map showing the log_2_ fold change of RLU produced by each CaM/CML13/14 relative to CML42 as the negative control. Each box on the heat map represents the mean of six technical replicates of a representative biological replicate (n = 4). An asterisk indicates a significant difference to CML42 for 3 out of 4 biological replicates (one-way ANOVA against CML42 with Sidak’s test for multiple comparisons, P<0.05). RLU, relative light units.

To further test if CML13, CML14, and CaM bind to the neck domains of myosin XIs within plant cells, a Cytoskeleton-Based Assay for Protein-Protein Interaction, CAPPI assay, was performed (Lv *et al*., 2017) (Fig 3). The CAPPI assay was utilized instead of the myosin tail domains as the localization of the myosin XI tails *in planta* was previously found to be diffuse in the cytoplasm with no particular site of accumulation (Avisar *et al*., 2009). CML13, CML14, and CaM-GFP fusions were co-expressed with microtubule associating-protein (MAP) 65-1 fused to RFP or to myosin XI-A, -H, -I, and -K neck domains in *N. benthamiana*. The MAP-65-1-RFP fusion was used as a negative control showing that CML13, CML14, and CaM did not localize to the microtubule network when transiently expressed *in planta* (Fig. 3). In contrast, when co-expressed with MAP65-1-XI neck domains, CML13 and CML14-GFP clearly re-localized to the cytoskeleton, likely associating with the microtubule network (Fig. 3). CaM-GFP on the other hand, did not appreciably re-localize to the cytoskeleton when co-expressed with MAP-65-1-XI-A or -H, but showed modest re-localization with microtubules when co-expressed with XI-I and -K MAP65-1 fusions (Fig. 3).

**Figure 3.**
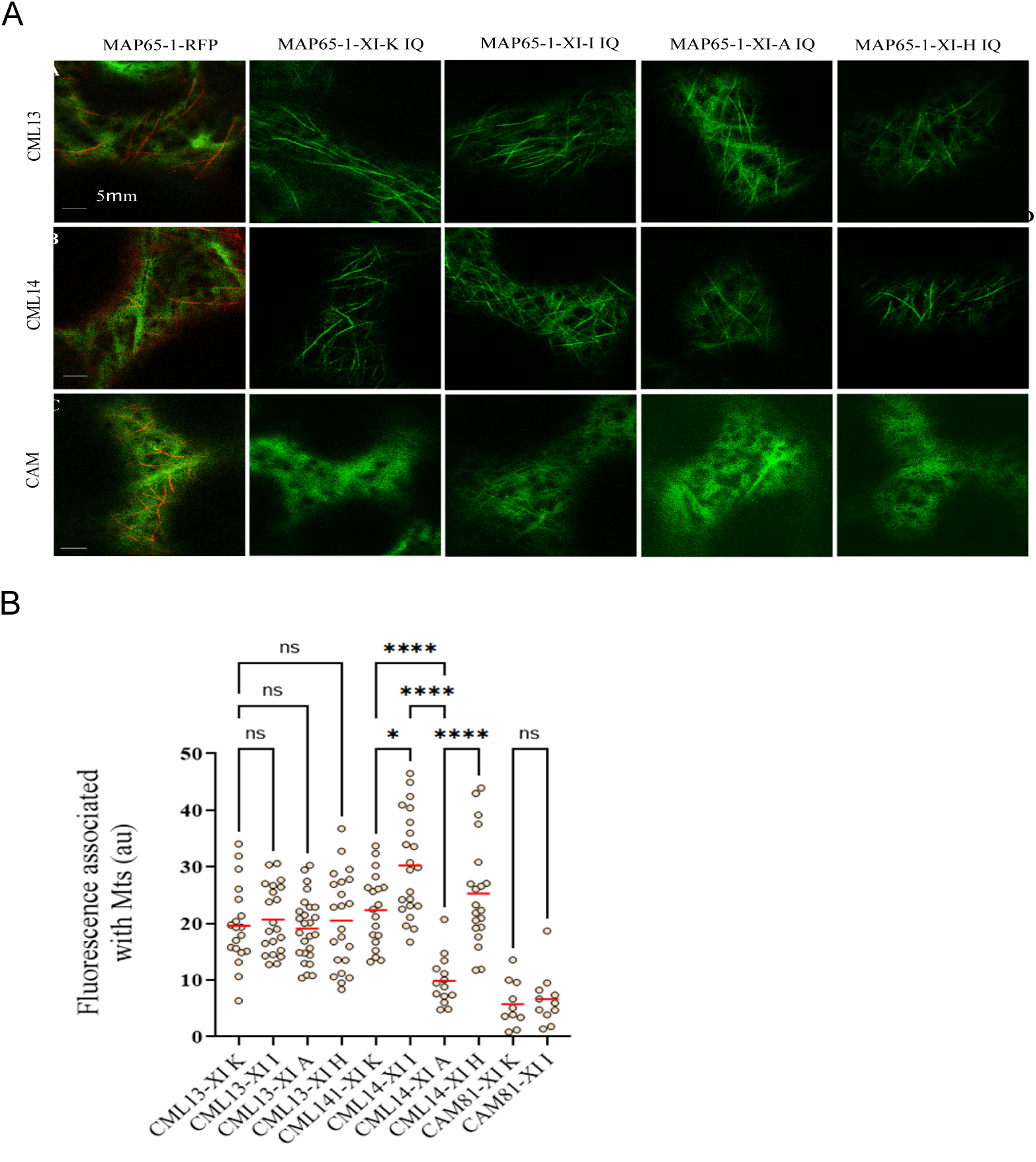
CAPPI analysis of Arabidopsis CaM/CML13/14 with MAP65-1 myosin XI IQ fusions. *N. benthamiana* leaves were co-infiltrated with *Agrobacterium* harboring a CML-GFP and MAP65-1-RFP or -myosin XI neck plasmid and were allowed to transiently express for 2 days. Microscopy was performed 48 h after agro-infiltration using a Leica SP8 confocal microscope and imaging was performed with using HyD detectors, HC PL APO CS 63x /1.2 water immersion objective (Leica, Wetzlar, Germany), an OPSL 488 laser for GFP excitation with 500–530 nm emission range, and an OPSL 552 laser for RFP with 565–640 nm emission light detection. A) MAP65-1-RFP was utilized as the negative control and highlighted the microtubule network within the *N. benthamiana* cells. MAP65-1-XI-A, MAP65-1-XI-H, MAP65-1-XI-I, MAP65-1-XI-K were co-expressed with CML13/14/CaM-GFP fusions to assess their re-localization *in planta*. B) A line was drawn with the ImageJ free-hand tool along and near the microtubules (MTs) to measure MT co-localization and subtract the background, respectively. The points on the graph represent a biological replicate of the mean line intensity along the MTs subtracted by the background fluorescence in the cell. Statistical analysis was by one-way ANOVA, *<P0.05, **P<0.01, *** P<0.001, ****P<0.0001. All microscopy images are at the same magnification as indicated in panel A.

### CML13, CML14, and CaM Have Different Specificities for Myosin XI IQ Motifs

As myosin XIs have six IQ motifs, we explored whether they exhibited specificity for the MLCs identified. Myosins XI-A, XI-H, XI-I, and XI-K were chosen as representative members because they span the phylogenetic tree (Fig. 1B) and displayed strong signals in the SL assays with the full neck domains (Fig. 2B). We found that almost all IQ domains interacted with CML13 and CML14 from each of the myosins tested. CaM did not interact with IQ2 of XI-A, H, I, or K, IQ4 from XI-A, H, or I, and IQ1 of XI-I. Myosin XI-H IQ6 was the only IQ6 motif that interacted with MLCs and interacted with CML13, CML14, and CaM in the SL assay.

These specificities were further investigated for myosin XI-A *in vitro* using the dansylated CaM/CML interaction assay (Alaimo *et al*., 2013). Single IQ domains were cloned into pET28a with the SUMO tag to increase the size and solubility of the IQ domain and recombinantly expressed in *E. coli.* SUMO-IQ myosin XI-A fusions were purified and tested for their interactions with dansylated-(D-)CaM, D-CML13, or D-CML14 in the presence or absence of calcium (Fig. 5). Each SUMO-IQ fusion was found to interact with D-CaM, D-CML13, and D-CML14 both in the presence and absence of calcium as the D-MLCs had significantly different fluorescent spectrums in the presence of the SUMO-IQ fusions (Fig. 5).

### CML13 and CML14 Have Different IQ Motif Contact Residues than CaM

To further investigate the *in planta* specificity of MLC-IQ interactions seen in Fig. 4, logo plots of the CML13/14 specific (non-CaM binding) and the non-specific (CaM binding) IQ motifs were made (Fig. 6). As expected, the IQ motif consensus residues are quite conserved (IQXXXRGXXXR) between the CaM binding and non-CaM binding IQ motifs except for the absence of the conserved Gly in the CaM binding motif sequence. Further, the Ala at positions 2, 3, and 15, and the F/Y at position 19 in the logo plots are conserved between the two motifs. The CaM binding motif possesses positively charged residues at positions 5, 8, and 17 that are largely absent in the non-CaM binding motif resulting in a markedly higher charge in these IQ domains. Conversely, the non-CaM binding motifs appear to possess more negatively charged amino acids than the CaM binding motifs. In place of the consensus Gly, the CaM binding IQ motif has a consensus Thr, or an Arg or Ser in some cases in this position which are large changes from the classic IQ consensus. To explore some of these differences further, and in an attempt to identify the residues important for CML13/14-IQ interaction, we made a series of point mutations in myosin XI-A IQ1 and IQ2 and tested these *in planta* using the SL assay (Fig. 7). Of the consensus residues, large changes to the IQXXXXXXXXRXXF residues had a marked reduction in the interaction of CML13, CML14, and CaM to IQ1. Mutations to the consensus IQXXXRGXXXR residues in IQ2 had the largest reduction of CML13 and CML14 interaction. Including the consensus residues, the IIQXXXLTXXXRXXF residues in IQ1 were all found to be important for MLC binding. Further, the residues SXEIQXXXRGXXXR in IQ2 were important for CML13 and CML14 interaction. It is hard to make generalized claims given that some of the mutations did not yield consistent changes in the interaction between IQ1 and IQ2. However, the mutation of the conserved Iso in the IQ to an Ala produced little change to CML13 or CML14 interaction while abolishing CaM interaction to IQ1. Further, mutations to IQ1 including T12F and F19G reduced the interaction of CML13 and CML14 to below background levels while retaining interaction with CaM. This implies that while CML13 and CML14 likely have overlapping IQ contact residues, CaM has unique contact/preferred residues within these IQ domains.

**Figure 4.**
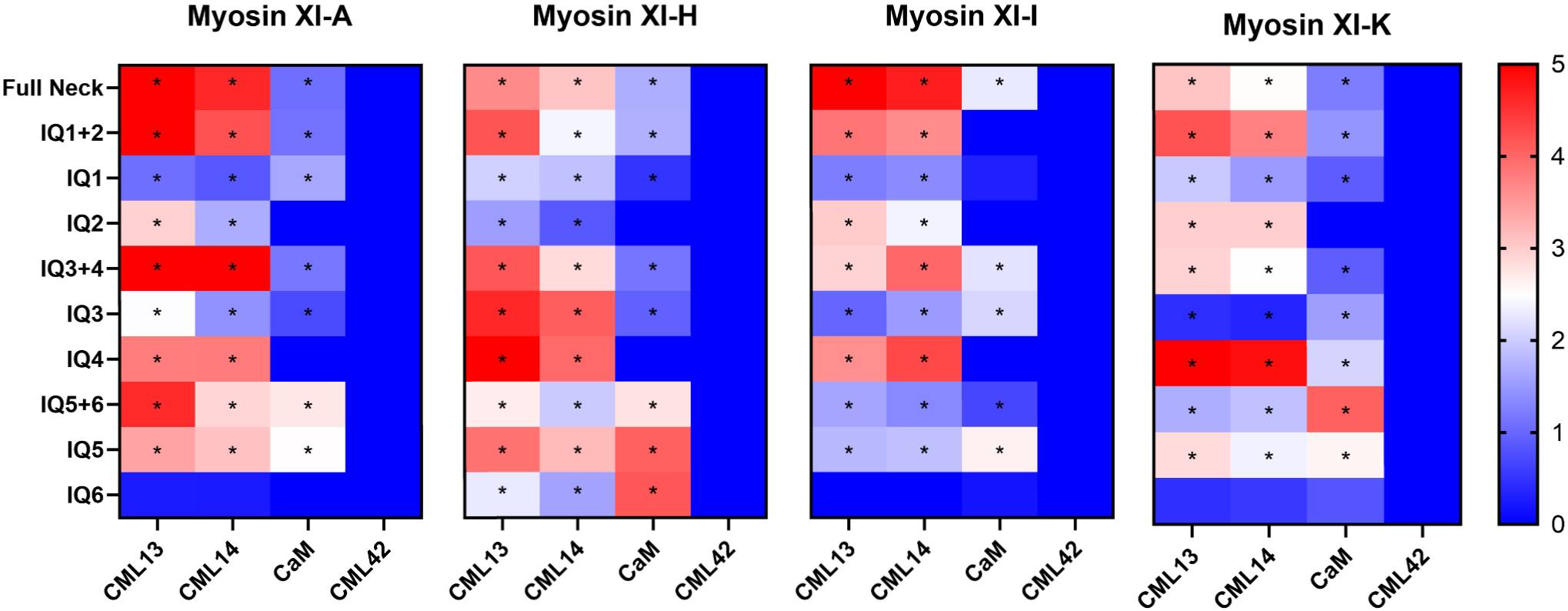
Split-luciferase protein interaction of Arabidopsis CaM, CML13, and CML14 with single and paired IQ domains of myosin XI-A, XI-H, XI-I, and XI-K *in planta*. *N. benthamiana* leaves were co-infiltrated with *Agrobacterium* harboring an NLuc-myosin and CLuc-CML plasmid and were allowed to transiently express these fusion proteins for 4 days. Heat maps depict the log_2_ fold change of RLU from CaM/CML13/14 relative to the RLU produced with CML42 and the same NLuc-myosin neck/IQ domain. The neck domains of myosins XI-A, -H, -I, and -K were delineated into their individual and pairs of IQ domains and tested for interaction with the putative MLCs as denoted in the figure. Each box on the heat map represents the mean of six technical replicates of a representative biological replicate (n = 4). An asterisk indicates a significant difference to CML42 for 3 out of 4 biological replicates (one-way ANOVA against CML42 with Sidak’s test for multiple comparisons, P<0.05). RLU, relative light units.

**Figure 5.**
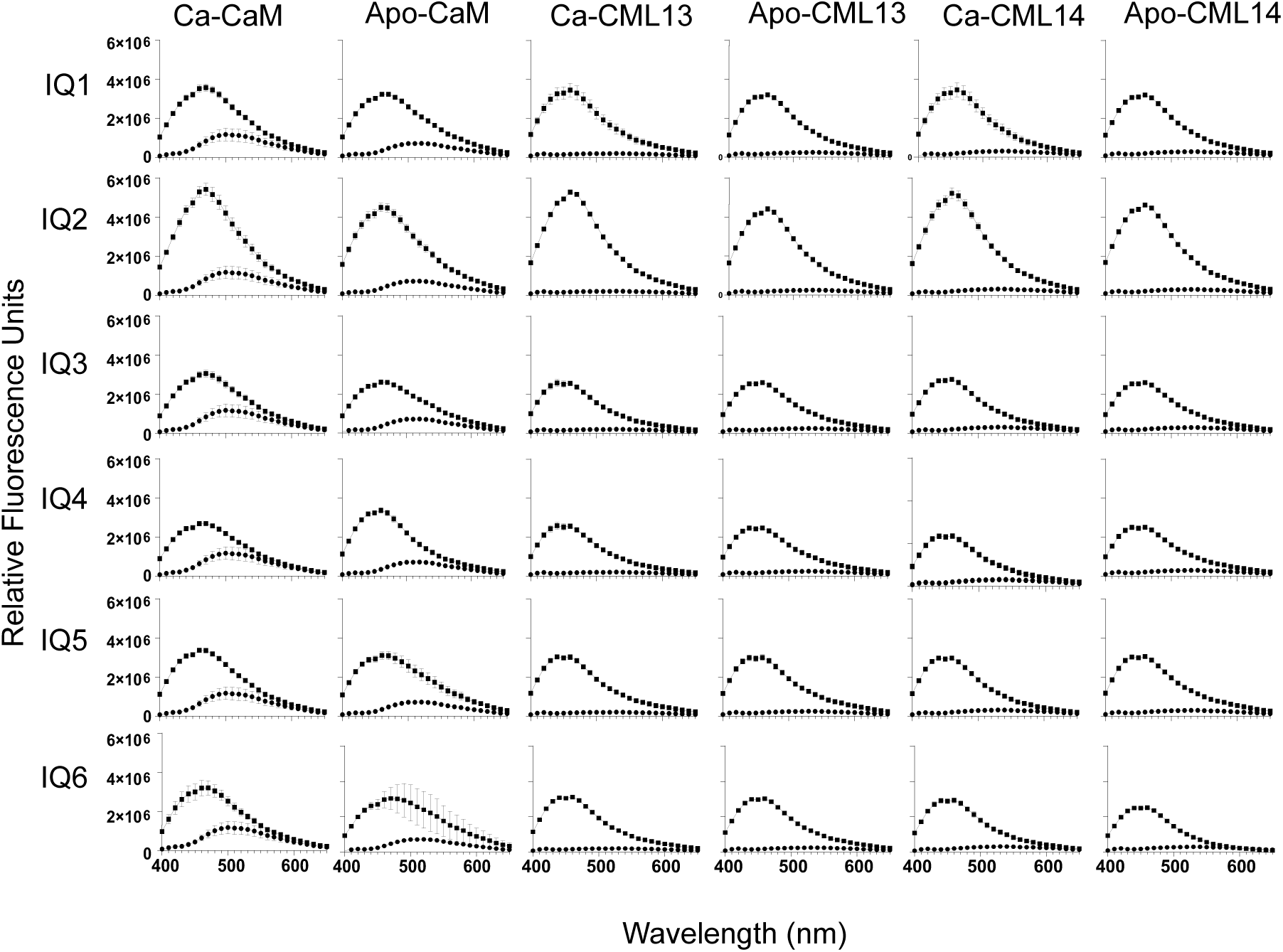
*In vitro* interaction of dansyl (D)-CaM, -CML13, or -CML14 with SUMO-IQ domain fusion proteins of the myosin XI-A IQ motifs. Dansyl fluorescence was measured over an emission wavelength (λ) window of 400 to 650 nm and an excitation wavelength of 360 nm. Samples of 3 µM D-CML13, D-CML14, or using 600 nM D-CaM, were separately tested for fluorescence alone (filled circle), or in the presence of SUMO-XIA-IQ fusion proteins (filled squares) supplemented with 1 mM CaCl_2_ (Ca-) or EGTA (Apo-). Fusion protein concentrations were used at a 10-fold molar excess. The intensity was measured in arbitrary relative fluorescence units (RFU). The fluorescence traces are the mean ±SD of three independent experiments performed in triplicate.

**Figure 6.**
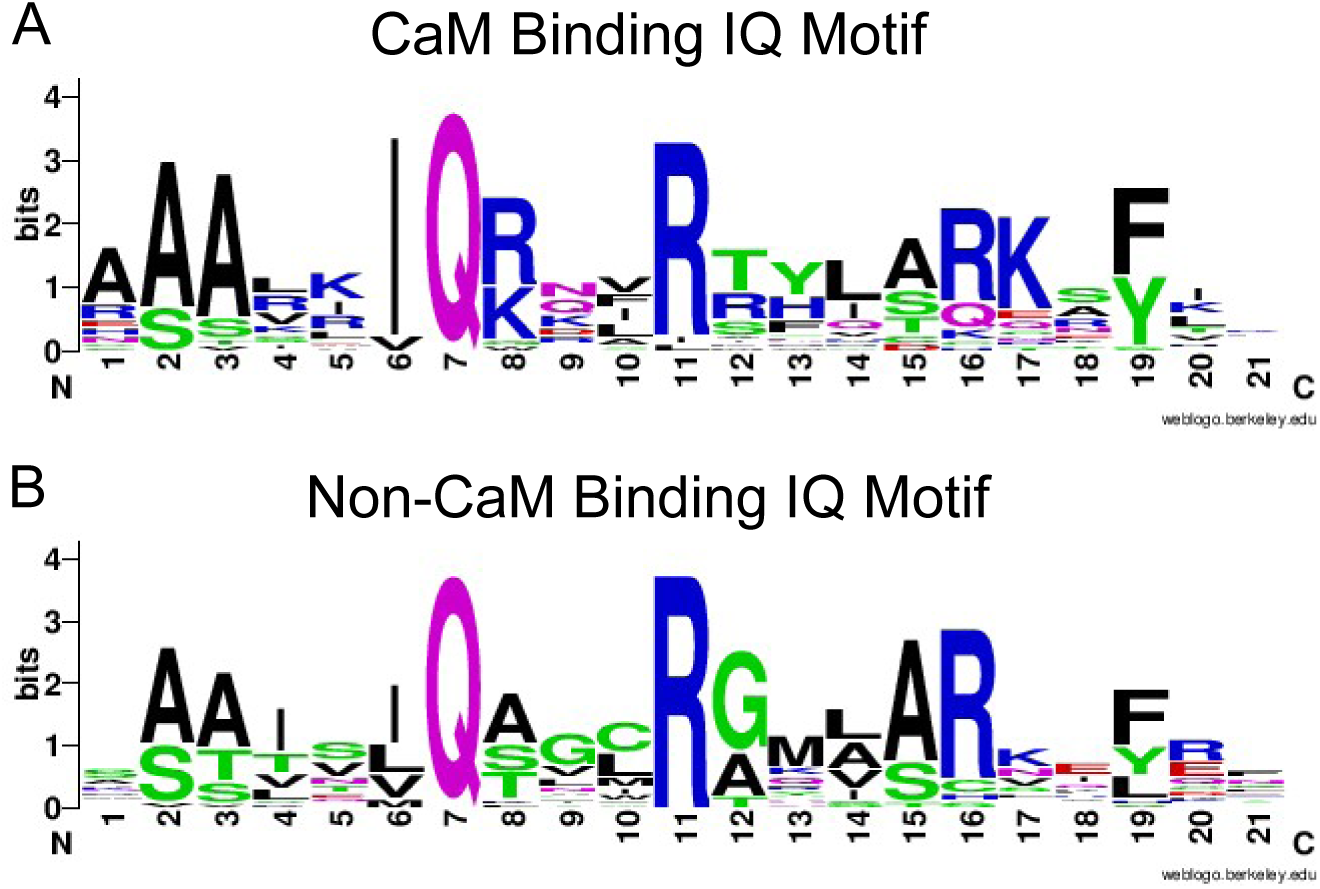
Logo plots of the CaM binding and non-CaM binding IQ motif sequences. Logo plots were generated from the myosin XI IQ motif sequences that interacted specifically in the *in planta* split-luciferase assay (Fig. 4) to CML13/14 (non-CaM binding) and the IQ sequences which bound to CML13/14/CaM (CaM binding). Logo plots were generated by web-logo and the size of each letter depicts the conservation at that residue within the motif (Crooks *et al*., 2004).

**Figure 7.**
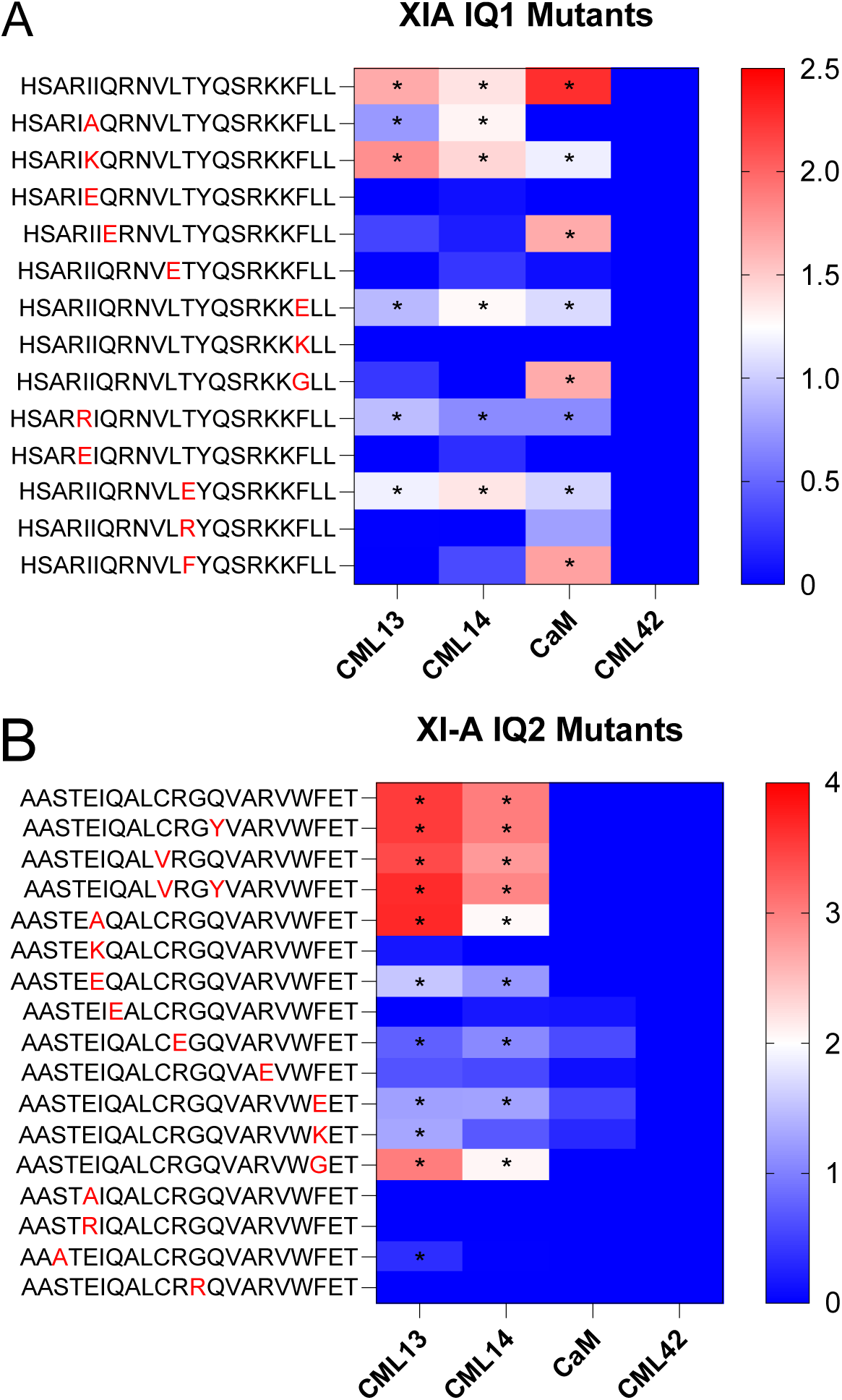
Split-luciferase protein interaction of Arabidopsis CaM, CML13, and CML14 with mutated IQ1 and IQ2 of myosin XI-A *in planta*. *N. benthamiana* leaves were co-infiltrated with *Agrobacterium* harboring an NLuc-XIA IQ and CLuc-CML plasmid and were allowed to transiently express these fusion proteins for four days. Heat maps depict the log_2_ fold change of RLU from CaM/CML13/14 relative to the RLU produced with CML42 and the same NLuc-myosin neck/IQ domain. The IQ1 and IQ2 domains of myosins XI-A were mutated as highlighted by the red residue in the IQ motif sequence and tested for interaction with the putative MLCs as denoted in the figure. Each box on the heat map represents the mean of six technical replicates of a representative biological replicate (n = 4). An asterisk indicates a significant difference to CML42 for three out of four biological replicates (one-way ANOVA against CML42 with Sidak’s test for multiple comparisons, P<0.05). RLU, relative light units.

### CML13, CML14, and CaM Function as Myosin Light Chains for Class XI Myosins

We next set out to evaluate the motility of myosin XI-A, -H, -I, and -K with different MLCs *in vitro.* Myosin XI-A, -H, -I, and -K constructs encoding the motor domains and native neck domains were expressed in High Five insect cells and purified by affinity chromatography. Unfortunately, myosin XI-A did not sufficiently express in insect cells and had to be excluded from our analysis. The actin sliding velocities of these myosin XIs were measured using an anti-Myc antibody-based version of the *in vitro* actin gliding assay (Ito *et al*., 2007). The velocities were tested in the presence of CaM, CML13, and CML14 alone and different combinations of MLCs as denoted in the figures (Fig. 8).

**Figure 8.**
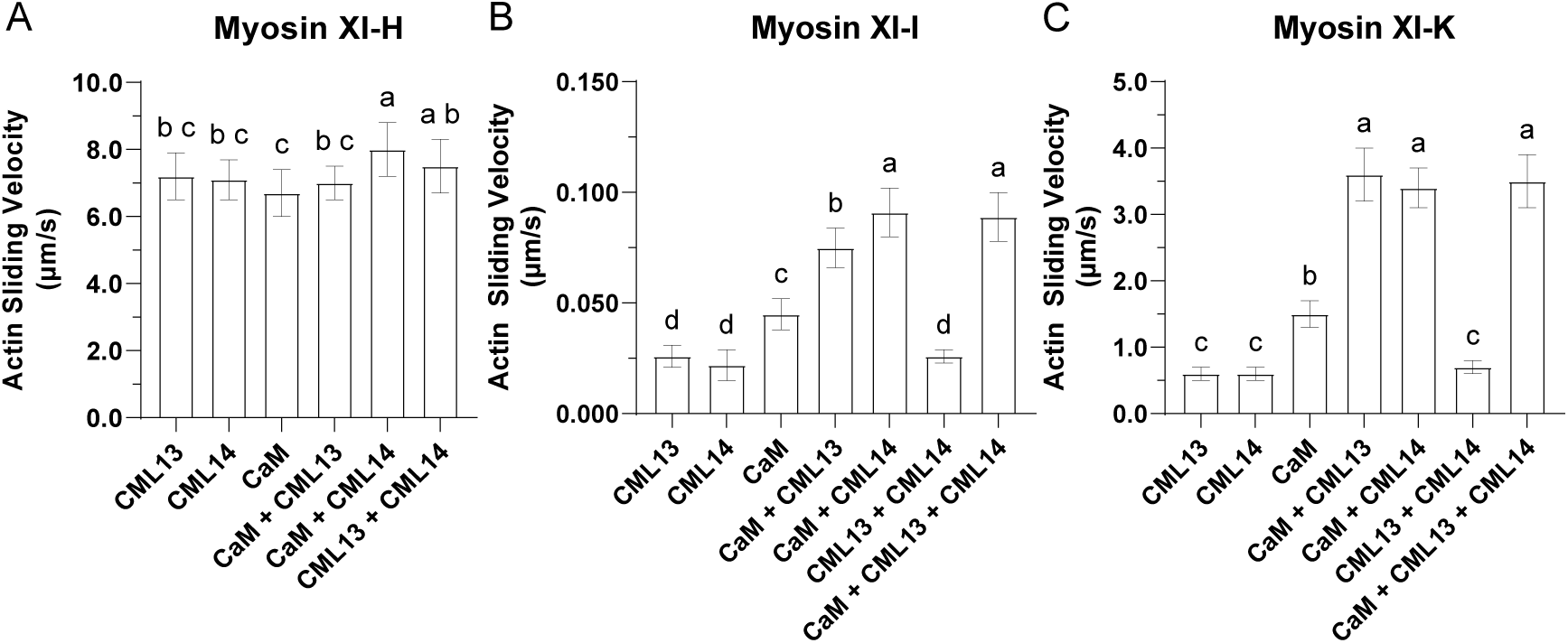
Actin sliding assays indicate that CaM, CML13, and CML14 can function as light chains for myosins XI-H, XI-I, and XI-K. Actin sliding velocity of myosin XI-H, -I, and -K in the presence of CaM, CML13, and CML14 alone or in different combinations of putative MLCs. The concentrations used for CaM, CML13, and CML14 were 30 µM each. Motility values are the mean±SD and were determined by measuring the displacements of actin filaments that were smoothly moving for distances >10 μm as described in the Materials and Methods. Unique letters above the means on the bar graph represent statistically significant differences between means (Two-way ANOVA with Dunnet’s test for multiple comparisons, P<0.05)

The *in vitro* velocities of XI-H with any single or combination of MLCs resulted in the maximal and smooth actin sliding (Fig. 8A). Myosin XI-I and XI-K motilities were increased by CaM more than either CML13 or CML14 alone and XI-I had maximal but not smooth actin sliding with the combination of CaM and CML14 while XI-K had maximal and smooth gliding with CaM and CML13/CML14 (Fig. 8B and C). Like the myosin VIIIs, myosin XIs can function with CaM and CML13 and/or CML14 as their MLCs *in vitro*.

### Partial Knockout of CML13 Phenocopies Myosin XI Mutants

A previous report found that the myosin XI triple mutant (XI-1, 2, K), *xi3KO*, has multiple physiological phenotypes, including reduced primary root growth compared to wild-type plants (Peremyslov *et al*., 2010). Moreover, we previously showed that silencing of *CML13* and *CML14* transcripts resulted in multiple phenotypes including the reduction or cessation of primary root growth (Symonds *et al*., 2024c). Here, we tested the myosin XI triple mutant along with myosin *XI-G*, *-H*, and *-I* single mutants and the myosin VIII quadruple mutant, *viii4KO,* alongside the *CML13* partial knockout, *cml13-1,* and the associated complementation lines for primary root growth (Talts *et al*., 2016; Ali *et al*., 2020) (Fig. 9). A confirmed knockout line of *CML14* was not available for analysis. After nine days of growth, only the *xi3KO* myosin mutant line showed a reduced primary root length that was rescued by the expression of XI-K-YFP in this background, as reported previously (Fig. 9A) (Peremyslov *et al*., 2010). The *cml13-1* line also had a statistically reduced primary root length compared to wild-type plants and this phenotype was rescued by the expression of *CML13-GFP, cml13R 1* and *2*, in the *cml13-1* background (Fig. 9B). This correlative data supports the hypothesis that CML13 functions in concert with at least some members of the myosin XI family in regulating plant growth.

**Figure 9.**
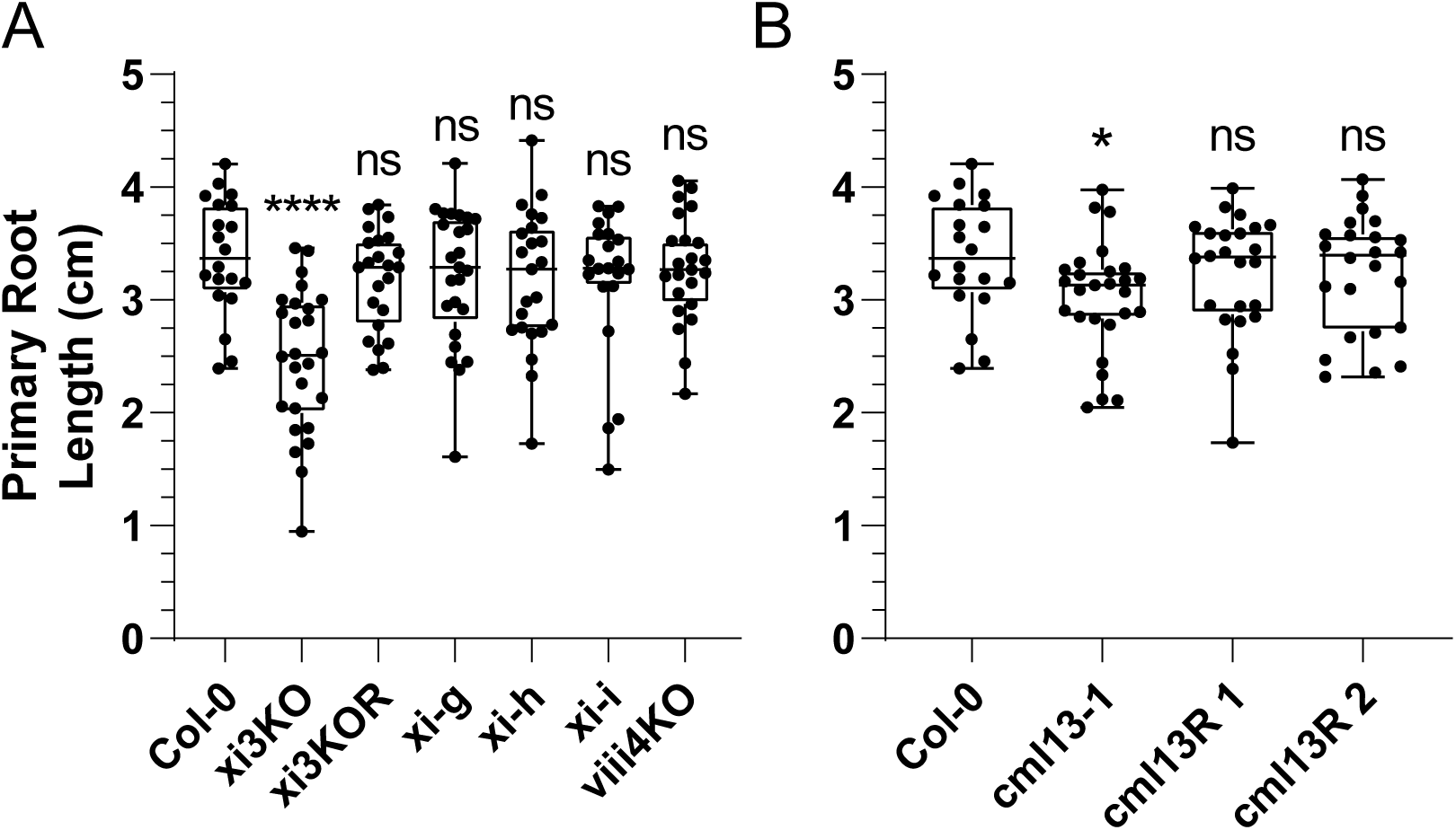
Primary root lengths of *CML13* and *myosin* mutants. Seeds were sown onto 0.5× MS agar plates and cold stratified in the dark at 4 °C for 48 h. Plates were then moved to the growth chamber and the seeds were allowed to germinate for four days before transplanting onto vertical plates for growth measurements and grown for another five days. The location of the primary root tip was marked daily and the final root length was measured using ImageJ. Box plots of (A) *myosin* and (B) *CML13* mutant lines show the mean with a horizontal bar and the outline of the boxes represents the 95% confidence interval, whiskers portray the range of the data acquired, and each point is a single biological replicate. Asterisks depict significant differences of mutant lines to Col-0 (one-way ANOVA with Tukey’s test for multiple comparisons, with 30–40 biological replicates, ns = not hsignificant, \**P*<0.05, \*\**P*<0.01, \*\*\**P*<0.001, \*\*\*\**P*<0.0001).

## Discussion

Despite the importance of myosin XI function in plant growth and development, the identity and properties of their myosin light chains (MLCs) has remained unknown for decades. Previous reports on myosin XIs have suggested that calmodulin (CaM) might serve as an MLC for myosin XIs *in vivo* (Yokota and Shimmen, 1994). However, it has been observed that CaM alone is insufficient for proper myosin XI function *in vitro* (Haraguchi *et al*., 2018). Our earlier study on Arabidopsis class VIII myosins revealed that CaM-like proteins, CML13 and CML14, along with CaM bind to the neck domains of all class VIII myosins and are sufficient for maximal *in vitro* motility of ATM1 and ATM2 (Teresinski *et al*., 2023; Symonds *et al*., 2024b). Based on this, we hypothesized and tested whether CML13 and CML14, in conjunction with CaM, serve as the MLCs for the larger class XI myosins. Multiple lines of evidence presented in this study support this hypothesis, including: (i) specific interaction and co-localization with the neck domains of all class XI myosins both *in planta* and *in vitro* (Fig. 2, 3, 5); (ii) an increase in *in vitro* myosin motility by CML13 and CML14 (Fig. 8); and (iii) the observation that the *cml13* mutant phenocopied the *xi3KO* primary root length phenotype (Fig. 9).

Using the SL interaction assay, we observed that CML13, CML14, and CaM interact with the neck domains of all class XI myosins *in planta* (Fig. 2). With the CAPPI (cytoskeleton-based assay for protein-protein interaction) assay we observed a slightly different result, CML13 relocalized in the presence of all myosin necks tested while CML14 only partially localized when co-expressed with myosin XI-A (Fig. 3). Also, CaM did not appear to relocalize with any of the myosin neck domains tested, however, this is likely due to the plethora of CaM targets in the cytosol competing for CaM-interaction *in planta* (Fig. 3). Further analysis using the SL assay revealed that CML13/14 and CaM do not have completely overlapping IQ interactors (Fig. 4). Specifically, CML13 and CML14 interacted with each IQ domain tested except for XI-A, XI-I, and XI-K IQ6, while CaM did not interact with IQ2 or IQ4 of the myosin XIs tested (Fig. 4). This pattern is reminiscent of the class VIII myosin neck domains, where IQ2 showed specificity for CML13/14 over CaM. The specificity of IQ-MLC interaction for CML13 or CML14 is further supported by motility assays, where maximal activity for myosin XI-I or XI-K required the combination of CaM and CML13/14, rather than any one MLC alone (Fig. 8B and C). However, similar to the myosin VIIIs and other IQ domain-containing proteins, the *in vitro* and *in planta* data did not always align. When tested *in vitro*, the specificity of IQ-MLC interaction disappeared, contradicting both the *in planta* data and the motility assays (Fig. 4, 5, 8). This discrepancy could be due to limitations of the *in vitro* assay, such as missing cellular components, or the truncation of the neck domain into individual IQs rather than pairs or a full neck domain may impact binding *in vitro*. The truncation was necessary to increase the solubility of these notoriously recalcitrant IQ domains (Homma *et al*., 2000; Bürstenbinder *et al*., 2017), but it may have reduced the structural context provided by flanking residues necessary for MLC specificity. Another contradiction involves the motility of myosin XI-H *in vitro* compared to the SL data and the other myosin XIs (Fig. 4, 8). While myosin XI-H IQ2 and IQ4 did not interact with CaM in the SL system, CaM alone produced approximately maximal XI-H motility *in vitro*. Notably, the combination of CaM and CML14 showed a statistically significant increase compared to CaM alone, although the differences between the means were only marginal (Fig. 8A).

It is a well-documented phenomenon that the myosin-based actin sliding velocity seen *in vitro* is linearly proportionate to the number of IQs in the neck domain (Haraguchi *et al*., 2018). Since myosin XIs possess six IQ domains, it is predicted that the motor-neck domains would be approximately six times faster *in vitro* than the motor alone (Haraguchi *et al*., 2018). Based on previously reported data for the motor activities, myosin XI-H and XI-K motor-neck domains produced approximately six times the actin sliding velocity of their motor domains alone (Haraguchi *et al*., 2018) (Fig. 8A and C). However, myosin XI-I, at the maximum, was only two times faster than the motor domain alone and this actin sliding was not smooth, suggesting that the neck domain was not fully occupied by CML13, CML14, and CaM (Fig. 8B). It is possible that under *in vitro* conditions, some factors are missing that allow CML13, CML14, and CaM to occupy the neck domain of myosin XI-I *in vivo*. For example, CML13 and CML14 have multiple reported phosphosites *in vivo* (Symonds *et al*., 2024c), which could be required for binding in this case, but this remains speculative. Furthermore, myosin XI-I contains a known phosphosite at the conserved Thr within IQ1, which appears to be unique among the class XI myosins (Mergner *et al*., 2020). It is currently unknown if this site is phosphorylated within HighFive insect cells and how this phosphorylation may impact motility, MLC binding, and XI-I function *in vivo*. However, it is also possible that myosin XI-I interacts with an as-of-yet unidentified MLC, perhaps another member of the CML family or related EF-hand protein. Myosin XI-I is functionally and phylogenetically unique, primarily functioning in maintaining nuclear shape (Tamura *et al*., 2013; Zhou *et al*., 2015). The *Zea mays* orthologue, Opaque1, was shown to play a role in asymmetric cell division through phragmoplast guidance, rather than actively participating in transport (Nan *et al*., 2023). This uniqueness may necessitate alternative regulatory mechanisms or proteins to control its motility compared to its other family members. Future work on myosin XIs and their light chains should investigate these hypotheses and whether CML13, CML14, and CaM are sufficient for maximal and smooth activity of the other class XI myosins.

Multiple IQ domains within the neck region of myosins allow MLCs to bind to and physically stabilize the neck, enabling it to act as a rigid lever arm that propagates the motile force. The IQ motif is generally described by the IQXXXRGXXXR consensus sequence, however, there is significant variation within these domains (Andrews *et al*., 2020). The typical CaM-binding IQ motif is thought to follow the consensus IQXXΦRGΦXXRXXΦ, where Φ represents a hydrophobic residue (Houdusse *et al*., 2006; Mori *et al*., 2008). Interestingly, the CaM-binding IQ motifs identified in this study—IQ1, IQ3, and IQ5 of myosin XI-A, XI-H, XI-I, and XI-K—also generally adhere to this consensus, with a substitution of Gly for Thr/Ser/Arg based on the logo plots (Fig. 4, 6). In contrast, the non-CaM binding IQ logo plot shows the consensus IQXXXRGXXXRXXΦ and lacks the conserved Φ residues flanking the Arg-Gly. To investigate whether these residues confer MLC-IQ specificity, we swapped the corresponding residues on myosin XI-A IQ2 with those of XI-A IQ1 (Fig. 7). These mutations, both individually and in combination, did not result in CaM binding to the mutated IQ2, indicating that these residues alone do not confer specificity (Fig. 7). Another potential differentiating factor between CaM and non-CaM binding IQ motifs is the abundance of positively charged amino acids in the CaM binding IQs. Previous reports suggest that positively charged residues enhance CaM affinity by forming electrostatic interactions with the negatively charged CaM protein (André *et al*., 2004). Indeed, we observed that mutations to Glu (E) led to a decrease in RLU produced with CaM, while CML13 and CML14 RLU signals showed little change (Fig. 7A). This implies that CaM is more sensitive to the charge of the binding domain than CML13 and CML14. Additionally, a series of point mutations revealed that residues IIQXXXLTXXXRXXF in IQ1 and SXEIQXXXRGXXXR in IQ2 are important for CaM and/or CML13 and CML14 binding, respectively (Fig. 7). The sequence variations among IQ domains and the contact residues between CML13/14-IQ domains remain open questions. While solving the CML13/14-IQ complex structure through X-ray crystallography or NMR solution structure would be the next step, these analyses were beyond the scope of this study. Furthermore, due to the variability of contact residues between CaM and IQ domains reported in the literature, this pursuit may not yield a generalized IQ motif that interacts with CML13 or CML14 (Houdusse *et al*., 2006; Mori *et al*., 2008; Shen *et al*., 2016). An additional challenge to future biochemical and structural analysis is the inherent insolubility of IQ domains.

The regulation of native myosin XI motility *in vivo* and *in vitro* by calcium has been previously studied (Yokota *et al*., 1999; Li *et al*., 2011). Earlier reports suggested that similar to mammalian class V myosins, calcium inhibits motility by dissociating CaM from IQ2 within the neck domain (Batters and Veigel, 2016). However, our data challenges this hypothesis. We found that CML13 and CML14 primarily occupy IQ2 instead of CaM *in planta* (Fig. 4). This observation aligns with what we noticed in Arabidopsis class VIII myosins, where CML13/14, being calcium-insensitive, likely do not play a role in calcium regulation (Vallone *et al*., 2016; Teresinski *et al*., 2023). We propose that CaM bound to IQ1 changes conformation, rather than dissociating, in the presence of calcium. This conformational change could decrease myosin motility with elevated calcium concentrations by affecting the actin binding/ATP hydrolysis rate of the motor domain. Our D-CaM ATM1 and ATM2 IQ1 interaction data supports this hypothesis, showing that CaM changes conformation while bound to the IQ domain *in vitro* (Symonds *et al*., 2024b). Additionally, crystallography data of CaM in complex with IQ1 of mouse myosin Va is consistent with this hypothesis (Houdusse *et al*., 2006). The N-lobe of CaM markedly changes its orientation in the presence of calcium compared to its absence (Houdusse *et al*., 2006). Further, the dissociation hypothesis often required >100 µM calcium concentrations to observe CaM dissociation, which would likely never occur *in vivo* (Yokota *et al*., 1999; Manceva *et al*., 2007). The biophysical mechanism of calcium regulation in myosin XIs and other unconventional myosins warrants further investigation. A future study that pairs protein structural analysis with the enzymatic analysis of myosin motors could shed light on this topic.

The potential number of targets for CML13 and CML14 presents a significant hurdle in identifying specific physiological roles for these CMLs. CML13 and CML14 were shown to interact with members of the CAMTA, IQD, and myosin families, which in Arabidopsis would total 56 putative targets, each with multiple IQ domains (Teresinski *et al*., 2023; Hau *et al*., 2024; Symonds *et al*., 2024d). Thus, similar to CaM, the physiological roles of CML13/14 may be expansive, and it will be difficult to assign specific functions to these proteins *in vivo*. Both *CML13* and *CML14* are essential for plant growth and development as the silencing of either results in a pleiotropic phenotype that results in mortality (Symonds *et al*., 2024c). Further, they appear to have opposing roles in ABA signaling at the germination stage as silencing *CML13* increased, and *CML14* decreased, ABA sensitivity (Symonds *et al*., 2024c). The silencing of *CML13* and *CML14* was also found to phenocopy the shortened hypocotyl phenotype of myosin VIII quadruple KO, *viii4ko* (Symonds *et al*., 2024c). Additionally, the inhibition of *CML13* and *CML14* transcripts *in vivo* resulted in a stark decrease in primary root length, reminiscent of the myosin XI triple KO (XI-1, 2, K) root growth phenotype (Peremyslov *et al*., 2010; Symonds *et al*., 2024c). Further, in this report, we showed that the partial knockout of *CML13* phenocopies the myosin XI triple mutant primary root length phenotype and was rescuable by the expression of *CML13-GFP* in the *cml13-1* mutant background (Fig. 9). Given that the silencing of *CML13* and *CML14* transcripts and the partial knockout of *CML13* displayed phenotypes similar to the myosin XI-1, 2, K triple mutant, we speculate that CML13 and CML14 participate in myosin XI-1, 2, and K’s functions during primary root growth. Myosin XIs, including -1, -2, and -K, participate in a myriad of other physiological functions, and CML13 and CML14’s roles in those processes should be investigated in future research.

In our previous report we speculated that like class VIII myosins, class XI myosins utilize CML13 and CML14 along with CaM as MLCs. This report confirms this hypothesis through multiple lines of experimental evidence *in vitro*, *in planta,* and *in vivo.* Confirmation of CML13/14 as MLCs for myosin VIII and XI classes answers a decades-old question of the identity of some MLCs in Arabidopsis (Vahey *et al*., 1982; Ma and Yen, 1989). These findings should accelerate myosin research, especially regarding the regulation of myosin function *in vivo*. Among the remaining unanswered questions to be addressed with future research include; how IQ-MLC specificity is achieved, whether CML13, CML14, or IQ domain phosphorylation has an impact on MLC binding or myosin activity, and whether myosin XI-I and other family members use another currently unknown MLC *in vivo*.

## Materials and Methods

### Plant material and growth conditions

*Nicotiana benthamiana* and *Arabidopsis thaliana* (Col-0) seeds were sown in Sunshine mix #1 (Sun Gro Horticulture Canada Ltd) and transferred to a growth chamber under short-day (SD) or long-day (LD) conditions, respectively (Conviron MTR30; SD, 12 h photoperiod; LD, 18 h photoperiod, 22 °C, ∼150 μmol m^−2^ s^−1^). For *N. benthamiana*, 14-day-old seedlings were transplanted to independent 10 cm pots and returned to the growth chamber. All pots were sub-irrigated as needed, with N-P-K fertilizer (20-20-20, 1 g L^−1^) applied every other week. For fluorescence microscopy assays, *N. benthamiana* growth conditions were as previously described (Belausov *et al*., 2023). For primary root length assays, Arabidopsis seeds were sown onto 0.5× Murashige and Skoog (MS) agar plates before cold stratifying in the dark at 4 °C for 48 h. Plates were then moved to the growth chamber and seedlings were allowed to establish for 4 d before transplanting onto 0.5x MS agar media and grown vertically for another 5 days before measuring the root lengths with ImageJ (Rueden *et al*., 2017). The *CML13* and *CML14* lines were produced previously (Hau *et al*., 2024; Symonds *et al*., 2024c). The Myosin VIII 4KO line, hereafter referred to as myosin *viii4KO*, was previously described (Talts *et al*., 2016). The myosin XI mutants, *xi3KO* and *xi3KOR*, were also generated previously (Talts *et al*., 2016).

### Plasmid constructs and recombinant protein expression

For PCR and cloning, oligonucleotide primers used are listed in Supplementary Table S1. See Supplementary Table S2 for a description of plasmid constructs used in this study and the corresponding regions of proteins encoded by the respective cDNAs, and respective locus identifiers. The pCambia1300-C-Luciferase (CLuc) vectors containing the CaM or CML cDNA sequences were constructed previously (Teresinski *et al*., 2023). cDNAs encoding the full region or truncations of the Arabidopsis myosin class XI neck domains were cloned upstream of the N-terminus of firefly luciferase in the pCambia1300-N-Luciferase (NLuc) binary vector.

Plasmids encoding *CML13*, *CML14*, and *CaM* fused to *GFP* were previously described (Symonds *et al*., 2024b). Either *RFP mCherry* or *myosins IQ domain* (XI-A, XI-H, XI-I, and XI-K) were fused at the C terminus of *MAP65-1* sequence. The cDNA of *MAP65-1* with BsaI adapters lacking stop codon and myosin IQ domains constructs containing stop codon with BsaI adapters, were synthesized by Twist Bioscience (Supplementary Table S3). To prepare *MAP65-1 + IQ domain of myosins* (XI-A, XI-H, XI-I, XI-K) constructs, we used *35S* promoter and *OCS* (*octopine synthase*)-terminator sequences fetched from pART7 vector. We used Golden Gate cloning system to prepare fusion constructs (Engler *et al*., 2014). All level-0 plasmids were mixed with backbone binary plasmid (pICH47742) to create a level-1 plasmid by cut and ligate reaction using BsaI restriction enzyme and T4 DNA ligase from NEB (New England Biosciences). All plasmids were confirmed by restriction digestion and DNA sequencing. Agrobacterium infiltration was performed as previously described (Belausov *et al*., 2023) with some modifications. We used infiltration buffer containing 10 mM MgCl_2_, 10 mM MES pH 5.6, 150 µM acetosyringone for agroinfiltration and incubated the resuspended culture for two hours at 28°C at 50 RPM speed before infiltration.

For recombinant protein expression, the cDNA sequences of interest were subcloned into expression vectors (Supplementary Table S2). cDNAs encoding the IQ domains of *myosin XI-A* (At1g04600) were cloned into the pET28-SUMO vector for expression in *Escherichia coli* strain BL21 (DE3) CPRIL (Novagen). pET28-SUMO encodes SUMO, the small ubiquitin moiety, that has previously been successful at solubilizing IQ domain-containing and recalcitrant proteins (Damo *et al*., 2013; Symonds *et al*., 2024a). Recombinant proteins corresponding to the SUMO-IQ fusion of myosin XI-A were purified using Profinity Ni-IMAC column (BioRad) as per the manufacturer’s instructions. Recombinant CaM, CML13, and CML14 were expressed and purified as described (Teresinski *et al*., 2023).

For the myosin *in vitro* sliding assays, the myosin XI constructs encode the motor domain and native neck regions (six IQ motifs). Full-length cDNAs of *myosin XI-A, XI-H* (At4G28710), *XI-I* (At4g33200), and *XI-K* (At5g20490), were provided by Dr. Motoki Tominaga, respectively. A baculovirus transfer vector for each myosin was generated using PCR (Supplementary Table S1). PCR products were ligated into the NcoI–AgeI restriction sites of pFastBac MD (Ito *et al*., 2009). The resulting constructs, pFastBac myosin XI, encode N-terminal amino acids (MDYKDDDDKRS) containing the FLAG tag (DYKDDDDK), amino acid residues 1-883, 1-890, 1-889, and 1-884 of *XI-A, XI-H, XI-I,* and *XI-K,* respectively, and C-terminal amino acids (GGGEQKLISEEDLHHHHHHHHSRMDEKTTGWRGGHVVEGLAGELEQLRARLEHHPQGQRE PSR) containing a flexible linker (GGG), a Myc-epitope sequence (EQKLISEEDL), a His tag (HHHHHHHH), and a streptavidin-binding peptide (SBP) tag (MDEKTTGWRGGHVVEGLAGELEQLRARLEHHPQGQREP). Myosin XIs were expressed using a baculovirus expression system in HighFive™ insect cells. They were purified using nickel-affinity and FLAG-affinity resins as previously described (Haraguchi *et al*., 2022).

### Split-luciferase assay

The split-luciferase (SL) protein-protein interaction assays were performed as described (Chen *et al*., 2008; Teresinski *et al*., 2023). In brief, full cDNAs encoding ‘bait’ proteins (*CaM* and *CMLs*) were cloned into the CLuc binary vector for expression as fusion proteins in-frame with the C-terminal domain of firefly luciferase. The cDNAs encoding ‘prey’ proteins (regions of *myosin XIs* as indicated in the figures) were cloned into the NLuc binary vector for expression as fusion proteins in-frame with the N-terminal domain of firefly luciferase. Transformation and growth of *Agrobacterium tumefaciens* strain GV3101 were performed as described (Teresinski *et al*., 2023). Six-week-old *N. benthamiana* leaves were infiltrated by *A. tumefaciens* strain GV3101 carrying bait and prey constructs as indicated in the figures. Leaf discs were taken after four days, incubated with 100 μl of water containing 1 mM D-luciferin in a 96-well plate for 10–15 min, and luminescence was subsequently captured with the SpectraMax Paradigm multimode detection platform (Molecular Devices Inc.).

### Microscopy and image analysis

For the live cell imaging, a Leica SP8 confocal microscope was used. Imaging was performed using HyD detectors, HC PL APO CS 63x /1.2 water immersion objective (Leica, Wetzlar, Germany), an OPSL 488 laser for GFP excitation with 500–530 nm emission range, and an OPSL 552 laser for RFP with 565–640 nm emission light detection. The fluorescence associated with microtubules (MTs) was measured by the ImageJ using the line analysis tool (National Institutes of Health; USA http://imagej.nih.gov/ij). A line was drawn with the free-hand tool along the MTs and another line was drawn near the MTs for reference to remove the background noise. Florescence associated with MTs was calculated by subtracting the mean line intensity of the background from that of MTs. Graphs and statistics analysis were performed by GraphPad Prism 10 using One-way ANOVA.

### Steady-state dansyl fluorescence spectroscopy

Recombinant CaM and CMLs were covalently labelled using dansyl fluoride as described (Alaimo *et al*., 2013). Samples of 600 nM dansylated CaM (D)-CaM or 3 µM CMLs were incubated in TBS with 1 mM CaCl_2_ or 1 mM EGTA with or without a 10 molar excess of SUMO-IQ fusion, for 30 min at room temperature. Fluorescence spectra were collected using an excitation of 360 nm and emission wavelengths of 400–650 nm with 10 nm increments at 25 °C with a SpectaMax Paradigm (Molecular Devices) instrument. SUMO-IQ fusions were measured without the D-MLCs and this background fluorescence was subtracted from the spectrum of SUMO-IQ and D-MLC.

### Actin gliding assays

Actin sliding velocity was measured using an anti-Myc antibody-based version of the *in vitro* actin gliding assay as described (Ito *et al*., 2007). The velocity of actin filaments was measured in either 25 mM KCl (for myosin XI-K and XI-I) or 150 mM KCl (for myosin XI-H), 4 mM MgCl2, 25 mM HEPES-KOH (pH 7.4), 1 mM EGTA, 3 mM ATP, 10 mM DTT, and an oxygen scavenger system (120 μg/mL glucose oxidase, 12.8 mM glucose, and 20 μg ml^−1^ catalase) at 25 °C. CaM, CML13, and CML14 alone or in different combinations of putative MLCs were added to the assay buffer. Average sliding velocities were determined by measuring the displacements of actin filaments that were smoothly moving for distances >10 μm using ImageJ (Rueden *et al*., 2017) with the MTrackJ plugin (Meijering *et al*., 2012).

## Author contributions

WS, KS, LD, KI, ES, and MT designed the project, KS, LD, VD, EB, LP, and TH performed experiments and collected data. All authors contributed to data interpretation and manuscript writing.

## Conflict of interest

The authors have no conflicts to declare.

## Data Availability Statement

All data supporting the findings of this study are available within the paper and its supplementary materials published online.

## Funding

Funding was provided by a Natural Sciences and Engineering Council (NSERC) Discovery grant (2018-04928 to WS), the Israel Science Foundation (grant 626/22 to ES), by Grants-in-Aid for Scientific Research from the Japan Society for the Promotion of Science (JP 20001009, JP 23770060, and JP 23K05808 to MT; JP 17K07436, JP 26440131, JP 21570159, JP20K06583, 2JP 22H04833 and JP 23K05710 to KI; JP 22K20623 and JP 24K09482 to TH), and a grant from the Hamaguchi Foundation for the Advancement of Biochemistry to TH.

## Supplemental Tables

Supplemental Table S1. List of primers used in this study

Supplemental Table S2. Description of plasmids used in this study

Supplemental Table S3. Golden Gate cloning components

## Notes

### Competing Interest Statement

The authors have declared no competing interest.

