## Supplemental Tables for "The Calmodulin-like proteins, CML13 and CML14 Function as Myosin Light Chains for the Class XI Myosins in *Arabidopsis*"

**Supplementary Table S1**. List of primers used in this study

| Primer name | Sequence (5'-3') | Plasmid |
|---|---|---|
| XI-1 F | ggccggatccATGcttgggaatgctgccagagtc | NLuc |
| XI-1 R | ggccgtcgacACCGGTGTCTCTTGCAGCC | NLuc |
| XI-2 F | ggccggatccATGgtcttgggaagatcagcaagc | NLuc |
| XI-2 F | ggccggatccATGCGAAGGACTGAGGTCTTGGG | NLuc |
| XI-2 R | ggccgtcgacatGCAGGCGAGCCAGGTATCC | NLuc |
| XI-2 R | ggccggtcgacCTTAAGCTTTCGAAGTTCTCC | NLuc |
| XI-A F | ggccggatccATGGCTCACCGAGCCGAAGTTCTTGG | NLuc, pET SUMO |
| XIA IQ1 R | ggccgtcgacTTCTGTTGAAGCGGCCTGCAAC | NLuc, pET SUMO |
| XIA IQ1D F | ggccggatccATGGCTCACCGAGCCGAAGTTCTTGGTCATTCAGCACGGATTGCACAGAGA | NLuc |
| XIA IQ1E F | ggccggatccATGGCTCACCGAGCCGAAGTTCTTGGTCATTCAGCACGGATTAAACAGAGA | NLuc |
| XIA IQ1F F | ggccggatccATGGCTCACCGAGCCGAAGTTCTTGGTCATTCAGCACGGATTGAACAGAGA | NLuc |
| XIA IQ1G F | ggccggatccATGGCTCACCGAGCCGAAGTTCTTGGTCATTCAGCACGGATTATAGAGAGAAAT | NLuc |
| XIA IQ1H F | ggccggatccATGGCTCACCGAGCCGAAGTTCTTGGTCATTCAGCACGGATTATACAGAGAAATGTCGAAACTTAT | NLuc |
| XIA IQ1I R | ggccgtcgacTTCTGTTGAAGCGGCCTGCAACAACAAAAACTTCTTTTCAGATTG | NLuc |
| XIA IQ1J R | ggccgtcgacTTCTGTTGAAGCGGCCTGCAACAACAATTCCTTCTT | NLuc |
| XIA IQ1K R | ggccgtcgacTTCTGTTGAAGCGGCCTGCAACAACAATTTCTTCTT | NLuc |
| XIA IQ1L R | ggccgtcgacTTCTGTTGAAGCGGCCTGCAACAACAAACCCTTCTT | NLuc |
| XIA IQ1M F | ggccggatccATGGCTCACCGAGCCGAAGTTCTTGGTCATTCAGCACGGGCTATACAG | NLuc |
| XIA IQ1N F | ggccggatccATGGCTCACCGAGCCGAAGTTCTTGGTCATTCAGCACGGCGTATACAG | NLuc |
| XIA IQ1O F | ggccggatccATGGCTCACCGAGCCGAAGTTCTTGGTCATTCAGCACGGGAAATACAG | NLuc |
| XIA IQ1P R | ggccgtcgacTTCTGTTGAAGCGGCCTGCAACAACAAAAACTTCTTGCGAGATTGATATTCAAGGAC | NLuc |
| XIA IQ1Q R | ggccgtcgacTTCTGTTGAAGCGGCCTGCAACAACAAAAACTTCTTGCGAGATTGATATCTAAGGAC | NLuc |
| XIA IQ1R R | ggccgtcgacTTCTGTTGAAGCGGCCTGCAACAACAAAAACTTCTTGCGAGATTGATAAAAAAGGAC | NLuc |
| XIA IQ2 F | ggccggatccatgAAGAAGTTTTTGTTGTTGC | NLuc, pET SUMO |
| XIA IQ2 R | ggccgtcgacCCTCAGAGAAGCCGCTTC | NLuc, pET SUMO |
| XIA IQ2A R | ggccgtcgacCCTCAGAGAAGCCGCTTCTCTTCGCATGGTCTCAAACCAAACACGTGCAACATATCCTCTACAC | NLuc |
| XIA IQ2B R | ggccgtcgacCCTCAGAGAAGCCGCTTCTCTTCGCATGGTCTCAAACCAAACACGTGCAACTTGTCCTCTCACCAAGGCTTG | NLuc |
| XIA IQ2C R | ggccgtcgacCCTCAGAGAAGCCGCTTCTCTTCGCATGGTCTCAAACCAAACACGTGCAACATATCCTCTCACCAAGGCTTG | NLuc |
| XIA IQ2D F | ggccggatccatgAAGAAGTTTTTGTTGTTGCAGGCCGCTTCAACAGAAGCTCAAGCCTTG | NLuc |
| XIA IQ2E F | ggccggatccatgAAGAAGTTTTTGTTGTTGCAGGCCGCTTCAACAGAAAAACAAGCCTTG | NLuc |
| XIA IQ2F F | ggccggatccatgAAGAAGTTTTTGTTGTTGCAGGCCGCTTCAACAGAAGAGCAAGCCTTG | NLuc |
| XIA IQ2G F | ggccggatccatgAAGAAGTTTTTGTTGTTGCAGGCCGCTTCAACAGAAATTGAAGCCTTGTGTAG | NLuc |
| XIA IQ2H F | ggccggatccatgAAGAAGTTTTTGTTGTTGCAGGCCGCTTCAACAGAAATTCAAGCCTTGTGTGAAGGACAAGTTGC | NLuc |
| XIA IQ2I R | ggccgtcgacCCTCAGAGAAGCCGCTTCTCTTCGCATGGTCTCAAACCAAACTTCTGCAACTTG | NLuc |
| XIA IQ2J R | ggccgtcgacCCTCAGAGAAGCCGCTTCTCTTCGCATGGTCTCTTCCCAAACACG | NLuc |
| XIA IQ2K R | ggccgtcgacCCTCAGAGAAGCCGCTTCTCTTCGCATGGTCTCTTTCCAAACACG | NLuc |
| XIA IQ2L R | ggccgtcgacCCTCAGAGAAGCCGCTTCTCTTCGCATGGTCTCAGCCCAAACACG | NLuc |
| XIA IQ2M F | ggccggatccATGAAGAAGTTTTTGTTGTTGCAGGCCGCTTCAACAGCAATTCAA | NLuc |
| XIA IQ2N F | ggccggatccATGAAGAAGTTTTTGTTGTTGCAGGCCGCTTCAACACGAATTCAA | NLuc |
| XIA IQ2O F | ggccggatccATGAAGAAGTTTTTGTTGTTGCAGGCCGCTGCAACAGAA | NLuc |
| XIA IQ2P F | ggccggatccATGAAGAAGTTTTTGTTGTTGCAGGCCGCTTCAACAGAAATTCAAGCCTTGTGTAGAGAACAAGTT | NLuc |
| XIA IQ2Q F | ggccggatccATGAAGAAGTTTTTGTTGTTGCAGGCCGCTTCAACAGAAATTCAAGCCTTGTGTAGAAGACAAGTT | NLuc |
| XIA IQ2R F | ggccggatccATGAAGAAGTTTTTGTTGTTGCAGGCCGCTTCAACAGAAATTCAAGCCTTGTGTAGATTTCAAGTT | NLuc |
| XIA IQ3 F | ggccggatccatgGTTTGGTTTGAGACCATGC | NLuc, pET SUMO |
| XIA IQ3 R | ggccgtcgacTGAACAAGCAGACGAGCAC | NLuc, pET SUMO |
| XIA IQ4 F | ggccggatccatgAATGCCTATAAAACCCTGTG | NLuc, pET SUMO |
| XIA IQ4 R | ggccgtcgacAATGATTGTCGCTCTTCTC | NLuc, pET SUMO |
| XIA IQ5 F | ggccggatccatgGAACTTCAGTTAAGAAAGAAG | NLuc, pET SUMO |
| XIA IQ5 R | ggccgtcgacAGTAATTGCTGCTTTCTTCG | NLuc, pET SUMO |
| XIA IQ6 F | ggccggatccatgCAGCGTTATGTGAGAACG | NLuc, pET SUMO |
| XI-A R | ggccgtcgacAGCAGCCATTTTAAGATTTCGC | NLuc |
| XI-C F | ggccggatccATGgaggttctaagtagtgctgc | NLuc |
| XI-C F | ggccggatccATGGAGGTTCTAAGTAGTGCTGC | NLuc |
| XI-C R | ggccgtcgacTGCTCCTGTTTCCCTTGCAGCC | NLuc |
| XI-C R | ggccggtcgacTGCAGCCATTTTGAGCTTCC | NLuc |
| XI-D F | ggccggatccATGGTTCTTGGGCATTCAGCAAGG | NLuc |
| XI-D R | ggccagatcttTTCCTTAGCAGCCATTTTAAGC | NLuc |
| XI-E F | ggccggatccATGATGGTTCTCAGTGCTGCTGCC | NLuc |
| XI-E R | ggccggtcgacTTCCCTTGAAGCCATTTTGAG | NLuc |
| XI-F F | ggccggatccATGaccgaggtcttggctggtgcc | NLuc |
| XI-F R | ggccgtcgaccttGGCTTCCTTTAGTGCCCC | NLuc |
| XI-G F | ggccgagctcATGTTAGGAAGAGCAGCCTGC | NLuc |
| XI-G R | ggccggtcgacGTCCCTCGCATCCGTCTTAAGC | NLuc |
| XI-H F | ggccgagctcATGttaggcagagcagccagcagg | NLuc |
| XI-H IQ1 F | ggccggatccatgTTAGGCAGAGCAGCCAGCAGaATCCAG | NLuc |
| XI-H IQ1 R | ggccgtcgacATTAGTTGCAACCTTGCG | NLuc |
| XI-H IQ2 F | ggccggatccatgAAAACTTTTCTTATGCTTCG | NLuc |
| XI-H IQ2 R | ggccgtcgacCTCCAAAACAGCTGCATCTC | NLuc |
| XI-H IQ3 F | ggccggatccatgCTTATCTTTGAGGGCTTGC | NLuc |
| XI-H IQ3 R | ggccgtcgacGGAAACAGCAGCAAAATATAATTCC | NLuc |
| XI-H IQ4 F | ggccggatccatgCTTACAAGGAATTATATTTTGC | NLuc |
| XI-H IQ4 R | ggccgtcgacCATAATTGCCGCCTTGTCCTGC | NLuc |
| XI-H IQ5 F | ggccggatccatgGGCAGGCTCCGGTTCCAAAGG | NLuc |
| XI-H IQ5 R | ggccgtcgacTGTAATTGCTGCTTTCTTGAG | NLuc |
| XI-H IQ6 F | ggccggatccatgTTGCACTACCAGAGGCTCAAG | NLuc |
| XI-H IQ6 R | ggccgtcgacCCGTTTCCTTTGCAGCCATC | NLuc |
| XI-H R | ggccgtcgaccttGGCAGCTTCAAGGACTCCC | NLuc |
| XI-I F | ggccggatccATGCGGGCTGAAGTCCTTGATGC | NLuc |
| XI-I IQ 1-2 R | ggccggtcgacCAAGACAGCTGCCGCCGCAT | NLuc |
| XI-I IQ 3-4 F | ggccggatccATGAATGCTTATGCCACCAGAAGGA | NLuc |
| XI-I IQ 3-4 R | ggccggtcgacTAGAGAAGCAGCTCGATGCT | NLuc |
| XI-I IQ 5-6 F | ggccggatccATGTTAAAGTTTTCACATCAGAAAGAGC | NLuc |
| XII IQ1 R | ggccgtcgacTGAAATTGCAGAAGCCC | NLuc |
| XII IQ2 F | ggccggatccatgCAGAACTTCATCTCTGCAC | NLuc |
| XII IQ3 R | ggccgtcgacTACAATGGCAGCTGATAC | NLuc |

| | | |
|---|---|---|
| XII IQ4 F | ggccggatccatgTGTGCATTTGTAAAACTTG | NLuc |
| XII IQ5 R | ggccgtcgacAGCAATAATAGATGACTG | NLuc |
| XII IQ6 F | ggccggatccatgTCAGCATTCAGGCACCGTC | NLuc |
| XI-I R | ggccgtcgacAGCTAATCGCAAAGCACCTGC | NLuc |
| XI-J F | ggccggatccATGCTCGGCGAATCAGCGAGG | NLuc |
| XI-J R | ggccggtcgacTTTTCTGTCAGCCTGCTTTGAC | NLuc |
| XIK F | ggccggatccatgGTTCTCGGAAATGCAGCTAGGAG | NLuc |
| XI-K IQ 1-2 R | ggccggtcgacCTTCACCGCTGCTGCCTG | NLuc |
| XI-K IQ 3-4 F | ggccggatccATGAACCTTTATGAGGAAATGCGAC | NLuc |
| XI-K IQ 3-4 R | ggccggtcgacAATAGTAGCAGCCTTCATTTGC | NLuc |
| XI-K IQ 5-6 F | ggccggatccATGAATGAGTTCAGGTTTAGAAAGC | NLuc |
| XIK IQ1 R | ggccgtcgacCACGATAGCAGCTCCTCG | NLuc |
| XIK IQ2 F | ggccggatccatgGAATTCCGTGCTCTACGAG | NLuc |
| XIK IQ3 R | ggccgtcgacTGTTATTGTTGAATGTC | NLuc |
| XIK IQ4 F | ggccggatccatgGAATCTTATTTAAGAATTAG | NLuc |
| XIK IQ5 R | ggccgtcgacAGAAAGGGCAGCTTTCTG | NLuc |
| XIK IQ6 F | ggccggatccatgTCTTACTACAAGCAACTC | NLuc |
| XIK R | ggccgtcgacTGTCCTCAATTCTTTTCGTG | NLuc |
| XIA 6IQ F | aaaccATGGCTGCTTCAGCC | pFastBac |
| XIA 6IQ R | tttaccggtTGTTTCTTTAGCAGCCATT | pFastBac |
| XIH 6IQ F | aaaccATGGCTTGTACTACAGTCAATG | pFastBac |
| XIH 6IQ R | tttaccggtCGTTTCCTTTGCAGCC | pFastBac |
| XII 6IQ F | aaaccATGGCAAATTGTCTT | pFastBac |
| XII 6IQ R | tttaccggtTGCTTCATTAGCAACCTG | pFastBac |
| XIK 6IQ F | aaaccATGGTTGGCCC | pFastBac |
| XIK 6IQ R | tttaccggtTGTGTCTCGTGCGG | pFastBac |
| | | pFastBac |

**Supplementary Table S2**. Description of plasmid constructs used in this study

| Construct Name | AGI | Amino Acids | Vectors |
|---|---|---|---|
| CaM81 | M80836 (Genbank Accession) | Full | Cluc, pET5a |
| CML13 | At1G12310 | Full | Cluc, pET30a |
| CML14 | At1G62820 | Full | Cluc, pET30a |
| CML42 | At4G20780 | Full | Cluc, pET5a |
| Myosin XI-1 neck | AT1G17580 | 739-868 | NLuc |
| Myosin XI-2 neck | AT5G43900 | 789-937 | NLuc |
| Myosin XI-A neck | AT1G04600 | 728-880 | NLuc |
| Myosin XI-B neck | AT1G04160 | 789-937 | NLuc |
| Myosin XI-C neck | AT1G08730 | 750-882 | NLuc |
| Myosin XI-D neck | AT2G33240 | 750-899 | NLuc |
| Myosin XI-E neck | AT1G54560 | 736-886 | NLuc |
| Myosin XI-F neck | AT2G31900 | 730-890 | NLuc |
| Myosin XI-G neck | AT2G20290 | 744-888 | NLuc |
| Myosin XI-H neck | AT4G28710 | 725-890 | NLuc |
| Myosin XI-I neck | AT4G33200 | 742-895 | NLuc |
| Myosin XI-J neck | AT3G58160 | 732-880 | NLuc |
| Myosin XI-K neck | AT5G20490 | 733-886 | NLuc |
| Myosin XI-A IQ1 | AT1G04600 | 728-763 | NLuc, pET SUM[illegible] |
| Myosin XI-A IQ2 | AT1G04600 | 752-788 | NLuc, pET SUM[illegible] |
| Myosin XI-A IQ3 | AT1G04600 | 775-811 | NLuc, pET SUM[illegible] |
| Myosin XI-A IQ4 | AT1G04600 | 800-836 | NLuc, pET SUM[illegible] |
| Myosin XI-A IQ5 | AT1G04600 | 824-859 | NLuc, pET SUM[illegible] |
| Myosin XI-A IQ6 | AT1G04600 | 848-880 | NLuc, pET SUM[illegible] |
| Myosin XI-A IQ1+2 | AT1G04600 | 728-788 | NLuc |
| Myosin XI-A IQ3+4 | AT1G04600 | 775-836 | NLuc |
| Myosin XI-A IQ5+6 | AT1G04600 | 824-880 | NLuc |
| Myosin XI-H IQ1 | AT4G28710 | 725-770 | NLuc |
| Myosin XI-H IQ2 | AT4G28710 | 759-795 | NLuc |
| Myosin XI-H IQ3 | AT4G28710 | 782-818 | NLuc |
| Myosin XI-H IQ4 | AT4G28710 | 808-843 | NLuc |
| Myosin XI-H IQ5 | AT4G28710 | 830-866 | NLuc |
| Myosin XI-H IQ6 | AT4G28710 | 855-890 | NLuc |
| Myosin XI-H IQ1+2 | AT4G28710 | 725-795 | NLuc |
| Myosin XI-H IQ3+4 | AT4G28710 | 782-843 | NLuc |
| Myosin XI-H IQ5+6 | AT4G28710 | 830-890 | NLuc |
| Myosin XI-I IQ1 | AT4G33200 | 742-767 | NLuc |
| Myosin XI-I IQ2 | AT4G33200 | 758-794 | NLuc |
| Myosin XI-I IQ3 | AT4G33200 | 781-817 | NLuc |
| Myosin XI-I IQ4 | AT4G33200 | 806-842 | NLuc |
| Myosin XI-I IQ5 | AT4G33200 | 829-865 | NLuc |
| Myosin XI-I IQ6 | AT4G33200 | 854-895 | NLuc |

| | | | |
|---|---|---|---|
| Myosin XI-I IQ1+2 | AT4G33200 | 742-794 | NLuc |
| Myosin XI-I IQ3+4 | AT4G33200 | 781-842 | NLuc |
| Myosin XI-I IQ5+6 | AT4G33200 | 829-895 | NLuc |
| Myosin XI-K IQ1 | AT5G20490 | 733-763 | NLuc |
| Myosin XI-K IQ2 | AT5G20490 | 753-788 | NLuc |
| Myosin XI-K IQ3 | AT5G20490 | 775-811 | NLuc |
| Myosin XI-K IQ4 | AT5G20490 | 800-836 | NLuc |
| Myosin XI-K IQ5 | AT5G20490 | 823-859 | NLuc |
| Myosin XI-K IQ6 | AT5G20490 | 848-875 | NLuc |
| Myosin XI-K IQ1+2 | AT5G20490 | 733-788 | NLuc |
| Myosin XI-K IQ3+4 | AT5G20490 | 775-836 | NLuc |
| Myosin XI-K IQ5+6 | AT5G20490 | 823-875 | NLuc |

**Supplemental Table 3.** Golden Gate Cloning Components

| **Constructs** | **Sequences with BsaI adapters** |
|---|---|
| MAP65-1 | **CGGTCTCAAATG**GCAGTTACAGATACTGAAAGTCCTCATCTTGGGGAAATTACTTGTGGTACCTTACTTGAGAAGTTGCAGGAAAT<br>CTGGGATGAAGTTGGTGAGAGTGATGATGAACGAGACAAACTGCTTCTTCAGATAGAGCAAGAGTGTCTTGACGTTTACAAGAGAAAA<br>GTCGAGCAGGCTGCGAAATCCCGAGCTGAGCTTCTTCAAACCTTGTCAGATGCTAATGCTGAACTTTCCAGCCTCACAATGTCTCTTG<br>GAGACAAAAGCTTAGTTGGCATTCCGGATAAGTCTTCAGGAACGATTAAAGAACAACTTGCTGCAATAGCACCGGCTCTTGAACAACT<br>GTGGCAACAGAAAGAGGAGAGAGTCCGAGAGTTCTCTGATGTACAATCACAGATTCAGAAGATATGTGGAGATATTGCTGGAGGTTTG<br>AGCAATGAGGTTCCTATAGTCGATGAGTCTGATTTGTCACTGAAGAAATTAGACGATTTCCAGAGCCAACTCCAAGAGCTCCAGAAAG<br>AGAAGAGTGACAGGCTGCGCAAGGTGTTAGAGTTTGTGAGTACTGTTCATGATCTATGTGCTGTTCTTGGTTTGGATTTCTTAAGCAC<br>CGTCACCGAAGTTCATCCGAGCTTAGATGAAGATACCAGTGTCCAGTCTAAGAGCATTAGCAATGAGACTCTTTCAAGGTTGGCTAAA<br>ACCGTCTTGACTCTTAAAGATGATAAGAAGCAAAGACTTCAAAAGCTTCAAGAGCTGGCTACTCAGCTAATTGACCTGTGGAATCTGA<br>TGGATACTCCTGATGAGGAAAGAGAGCTTTTTGATCATGTTACCTGTAACATTTCATCTTCAGTCGATGAGGTCACTGTGCCAGGTGC<br>TCTTGCACGTGATTTGATTGAGCAGGCTGAGGTGGAAGTTGATAGGCTTGACCAGCTGAAAGCTAGCCGAATGAAAGAAATTGCGTTC<br>AAGAAGCAATCTGAGCTTGAAGAGATATATGCTCGTGCCCATGTAGAAGTTAACCCGGAATCTGCTCGTGAGAGAATCATGTCGCTGA<br>TTGATTCTGGAAACGTTGAGCCTACTGAATTATTGGCAGACATGGATAGCCAGATATCAAAGGCTAAGGAAGAAGCATTTAGTAGAAA<br>AGATATATTGGACCGAGTCGAGAAATGGATGTCAGCTTGTGAGGAAGAGAGCTGGCTAGAAGACTACAATCGGGATCAGAACAGGTAC<br>AGCGCAAGCAGAGGTGCACACTTGAATCTCAAGAGAGCTGAGAAAGCTCGGATTCTGGTTAGCAAGATTCCTGCCATGGTTGACACAT<br>TAGTTGCCAAGACCCGGGCTTGGGAAGAAGAACACAGCATGTCCTTTGCCTACGATGGTGTTCCTCTGCTAGCTATGCTAGACGAGTA<br>CGGTATGCTTAGGCAAGAACGAGAAGAGGAGAAACGGAGGCTGAGGGAACAAAAGAAGGTTCAAGAACAGCCACACGTAGAGCAAGAA<br>TCTGCCTTTAGCACCAGGCCAAGCCCTGCAAGACCGGTCAGTGCTAAGAAAACGGTGGGGCCACGAGCTAACAACGGAGGAGCCAATG<br>GAACACATAACCGGCGTTTATCTTTGAATGCAAACCAGAATGGAAGCAGGTCTACTGCAAAAGAAGCAGGGAGAAGGGAGACTCTCAA<br>CAGGCCGGCTGCTCCTACAAACTACGTTGCCATTTCGAAAGAGGAAGCTGCTTCATCTCCAGTTTCTGGTGCTGCAGATCATCAAGTT<br>CCAGCTTCACCA**TTCGTGAGACCG** |
| XIA IQ domain | **AGGTCTCATTCGGG**GTTCTTGGTCATTCAGCACGGATTATACAGAGAAATGTCCTTACTTATCAATCTCGCAAGAAGTTTTTGTTG<br>TTGCAGGCCGCTTCAACAGAAATTCAAGCCTTGTGTAGAGGACAAGTTGCACGTGTTTGGTTTGAAACCATGCGAAGAGAAGCGGCTT<br>CTCTGAGGATTCAGAAGCAGGCACGGACTTATATTTGCCAGAATGCCTATAAAACCCTGTGCTCGTCTGCTTGTTCAATTCAGACAGG<br>GATGCGAGCAAAAGCTGCTCGAATTGAACTTCAGTTAAGAAAGAAGAGAAGAGCGACAATCATTATTCAAAGTCAAATCAGAAGATGT<br>TTGTGTCATCAGCGTTATGTGAGAACGAAGAAAGCAGCAATTACTACTCAATGTGGCTGGAGAGTGAAAGTTGCACGCCGAGAATTGC<br>GATAAA**GCTTTGAGACCA** |
| XIH IQ domain | **AGGTCTCATTCGGG**GTCTTAGGCAGAGCAGCCAGCAGGATCCAGAGGAAATTCCGTTCCTACTTGTCCCGCAAAACTTTTCTTATG<br>CTTCGCAAGGTTGCAACTAATATGCAGGCTGTGTGCAGAGGTCAACTTTCTCGGCTTATCTTTGAGGGCTTGCGCAGAGATGCAGCTG<br>TTTTGGAGATTCAGAGAGACATACGAATGCATCTTGCTCGGAAGTCTTACAAGGAATTATATTTTGCTGCTGTTTCCATTCAATTAGG<br>AATACGTGGTATGGCTTCCCGTGGCAGGCTCCGGTTCCAAAGGCAGGACAAGGCGGCAATTATGATTCAGAGTCATTGTCGCAAATTC<br>CTGGCCCAGTTGCACTACCAGAGGCTCAAGAAAGCAGCAATTACAACACAAAGTGCATGGAGAGCAAGGTTAGCCCGTAAAGAACTGC<br>GGTAAA**GCTTTGAGACCA** |
| XII IQ domain | **AGGTCTCATTCGGG**TCTGCACGGGCTTCTGCAATTTCAATTCAGGCATACTGTAGAGGATGCCTGTCTCGAAATGCTTATGCCACC<br>AGAAGGAATGCGGCGGCAGCTGTCTTGGTCCAAAAGCATGTGCGCAGGTGGCTGTCAAGATGTGCATTTGTAAAACTTGTATCAGCTG<br>CCATTGTATTACAGTCTTGCATCCGTGCTGACTCAACTCGCTTAAAGTTTTCACATCAGAAAGAGCATCGAGCTGCTTCTCTAATTCA<br>GGCTCATTGGAGAATCCATAAGTTTCGCTCAGCATTCAGGCACCGTCAGTCATCTATTATTGCTATTCAGTGTCGTTGGCGACAGAAG<br>CTTGCGAAGAGAGAGTTTAGATAAA**GCTTTGAGACCA** |
| XIK IQ domain | **AGGTCTCATTCGGG**AATGCAGCTAGGAGAATTCAAAGGCAAAGCCGTACTTTTATTGCGTGCAAAGAATTCCGTGCTCTACGAGGA<br>GCTGCTATCGTGTTACAGTCTAACTGCCGAGGCAAATTGGCATGCAACCTTTATGAGGAAATGCGACGCCAGGCAGCAGCGGTGAAGA<br>TTCAGAAGATTTTCAGGAGACATATTGCCAGAGAATCTTATTTAAGAATTAGACATTCAACAATAACAGTTCAAACAGCGCTAAGGGG<br>AATGGTTGCTCGCAATGAGTTCAGGTTTAGAAAGCAAATGAAGGCTGCTACTATTATCCAGGCTCGACTTCGAAGTCATCTCACACAT<br>TCTTACTACAAGCAACTCCAGAAAGCTGCCCTTTCTACACAATGTGGCTGGAGGAGCAGGGTTGCACGAAAAGAATTGAGGTAAA**GC<br>TTTGAGACCA** |
| pART_35S promoter | **TGGTCTCAGGAG**TGAGACTTTTCAACAAAGGATAATTTCGGGAAACCTCCTCGGATTCCATTGCCCAGCTATCTGTCACTTCATCG<br>AAAGGACAGTAGAAAAGGAAGGTGGCTCCTACAAATGCCATCATTGCGATAAAGGAAAGGCTATCATTCAAGATCTCTCTGCCGACAG<br>TGGTCCCAAAGATGGACCCCCACCCACGAGGAGCATCGTGGAAAAAGAAGACGTTCCAACCACGTCTTCAAAGCAAGTGGATTGATGT<br>GACATCTCCACTGACGTAAGGGATGACGCACAATCCCACTATCCTTCGCAAGACCCTTCCTCTATATAAGGAAGTTCATTTCATTTGG<br>AGAGGACA**AATGTGAGACCA** |
| pART_OCS terminator | **TGGTCTCAGCTT**CTGCTTTAATGAGATATGCGAGACGCCTATGATCGCATGATATTTGCTTTCAATTCTGTTGTGCACGTTGTAAA<br>AAACCTGAGCATGTGTAGCTCAGATCCTTACCGCCGGTTTCGGTTCATTCTAATGAATATATCACCCGTTACTATCGTATTTTTATGA<br>ATAATATTCTCCGTTCAATTTACTGATTGTACCCTACTACTTATATGTACAATATTAAAATGAAAACAATATATTGTGCTGAATAGGT<br>TTATAGCGACATCTATGATAGAGCGCCACAATAACAAACAATTGCGTTTTATTATTACAAATCCAATTTTAAAAAAAGCGGCAGAACC<br>GGTCAAACCTAAAAGACTGATTACATAAATCTTATTCAAATTTCAAAAGGCCCCAGGGGCTAGTATCTACGACACACCGAGCGGCGAA<br>CTAATAACGTTCACTGAAGGGAACTCCGGTTCCCCGCCGGCGCGCATGGGTGAGATTCCTTGAAGTTGAGTATTGGCCGTCCGCTCTA<br>CCGAAAGTTACGGGCACCATTCAACCCGGTCCAGCACGGCGGCCGGGTAACCGACTTGCTGCCCCGAGAATTATGCAGCATTTTTTTG<br>GTGTATGTGGGCCCCAAATGAAGTGCAGGTCAAACCTTGACAGTGACGACAAATCGTTGGGCGGGTCCAGGGCGAATTTTGCGACAAC<br>ATGTCGAGGCTCAGCAG**CGCTTGAGACCA** |
